# Oxidation of Four Monoterpenoid Indole Alkaloid Classes by Three Cytochrome P450 Monooxygenases from *Tabernaemontana litoralis*

**DOI:** 10.1101/2024.07.29.605674

**Authors:** Zhan Mai, Kyunghee Kim, Matthew Bailey Richardson, Daniel André Ramey Deschênes, Jorge Jonathan Oswaldo Garza-Garcia, Mohammadamin Shahsavarani, Jacob Owen Perley, Destiny Ichechi Njoku, Ghislain Deslongchamps, Vincenzo De Luca, Yang Qu

**Author notes:** Corresponding authors: Ghislain Deslongchamps, Vincenzo De Luca, Yang Qu.

## Abstract

Cytochrome P450 monooxygenases (CYPs) are well known for their ability to catalyze diverse oxidation reactions, playing a significant role in the biosynthesis of various natural products. In the realm of monoterpenoid indole alkaloids (MIAs), one of the largest groups of alkaloids in nature, CYPs are integral to reactions such as hydroxylation, epoxidation, ring opening, ring rearrangement, and aromatization, contributing to the extensive diversification of these compounds. In this study, we investigate the transcriptome, metabolome, and MIA biosynthesis in *Tabernaemontana litoralis* (milky way tree), a prolific producer of rare pseudoaspidosperma-type MIAs. Alongside known pseudoaspidosperma biosynthetic genes, we identify and characterize three new CYPs that facilitate regio- and stereospecific oxidation of four MIA skeletons: iboga, aspidosperma, pseudoaspidosperma, and quebrachamine. Notably, the tabersonine 14,15-β-epoxidase catalyzes the formation of pachysiphine, the stereoisomer of 14,15-α-epoxytabersonine (lochnericine) found in *Catharanthus roseus* (Madagascar periwinkle) roots. The pseudovincadifformine 18-hydroxylase is the first CYP identified to modify a pseudoaspidosperma skeleton. Additionally, we demonstrate that the enzyme responsible for C10-hydroxylation of the iboga MIA coronaridine also catalyzes the same reaction on voaphylline, which bears a quebrachamine skeleton. With the discovery of a new MIA, 11-hydroxypseudovincadifformine, this study provides a comprehensive understanding of MIA biosynthesis and diversification in *T. litoralis*, highlighting its potential for further exploration.

## Introduction

Monoterpenoid indole alkaloids (MIAs) are renowned for their medicinal properties, intricate structures, and complex biosynthesis at both molecular and cellular levels. Most MIAs originate from the single precursor strictosidine, which is formed by the condensation of tryptamine and the monoterpenoid glucoside secologanin. The deglucosylation of strictosidine produces a highly labile dialdehyde intermediate, strictosidine aglycone, which is then reduced by various cinnamyl alcohol dehydrogenase (CAD)-like reductases to yield several primary MIA skeletons, including yohimbine, heteroyohimbine, and corynanthe types (Qu *et al*., 2017; Qu, Easson, *et al*., 2018; Stavrinides *et al*., 2016; Kim *et al*., 2023; Schotte *et al*., 2023; Stander *et al*., 2023). This reduction represents the first major diversification event in MIA biosynthesis. A family of homologous cytochrome P450 monooxygenases (CYPs) further catalyzes the intramolecular cyclization of the corynanthe MIA, geissoschizine, leading to the formation of sarpagan, akuammiline, and strychnos skeletons, thus marking the second major diversification event in MIA biosynthesis (Wang *et al*., 2022; Dang *et al*., 2018; Qu, Easson, *et al*., 2018). The consecutive oxidation and reduction of the key stychnos MIA *O*-acetylstemmadenine forms a chain-opened, unstable intermediate dehydrosecodine (Fig. 1). This intermediate serves as a key precursor to iboga, aspidosperma, and pseudoaspidosperma MIAs through intramolecular 4+2 cycloadditions and concomitant oxidation/reduction steps, marking the third major diversification event in MIA biosynthesis (Qu, Safonova, *et al*., 2018; Caputi *et al*., 2018; Kamileen *et al*., 2022; Farrow *et al*., 2019; Eng *et al*., 2022). The three skeletons inherit C2-C16 bond from dehydrosecodine but differ at C16-C21 bond (iboga), C17-C20 bond (aspidosperma), and C17-C14 bond (pseudoaspidosperma). All above basic skeletons can be further decorated and modified by numerous oxidoreductases, transferases, hydrolases, and other enzymes, leading to over 3,000 reported structures, therefore making MIA one of the largest alkaloid groups in nature.

**Figure 1.**
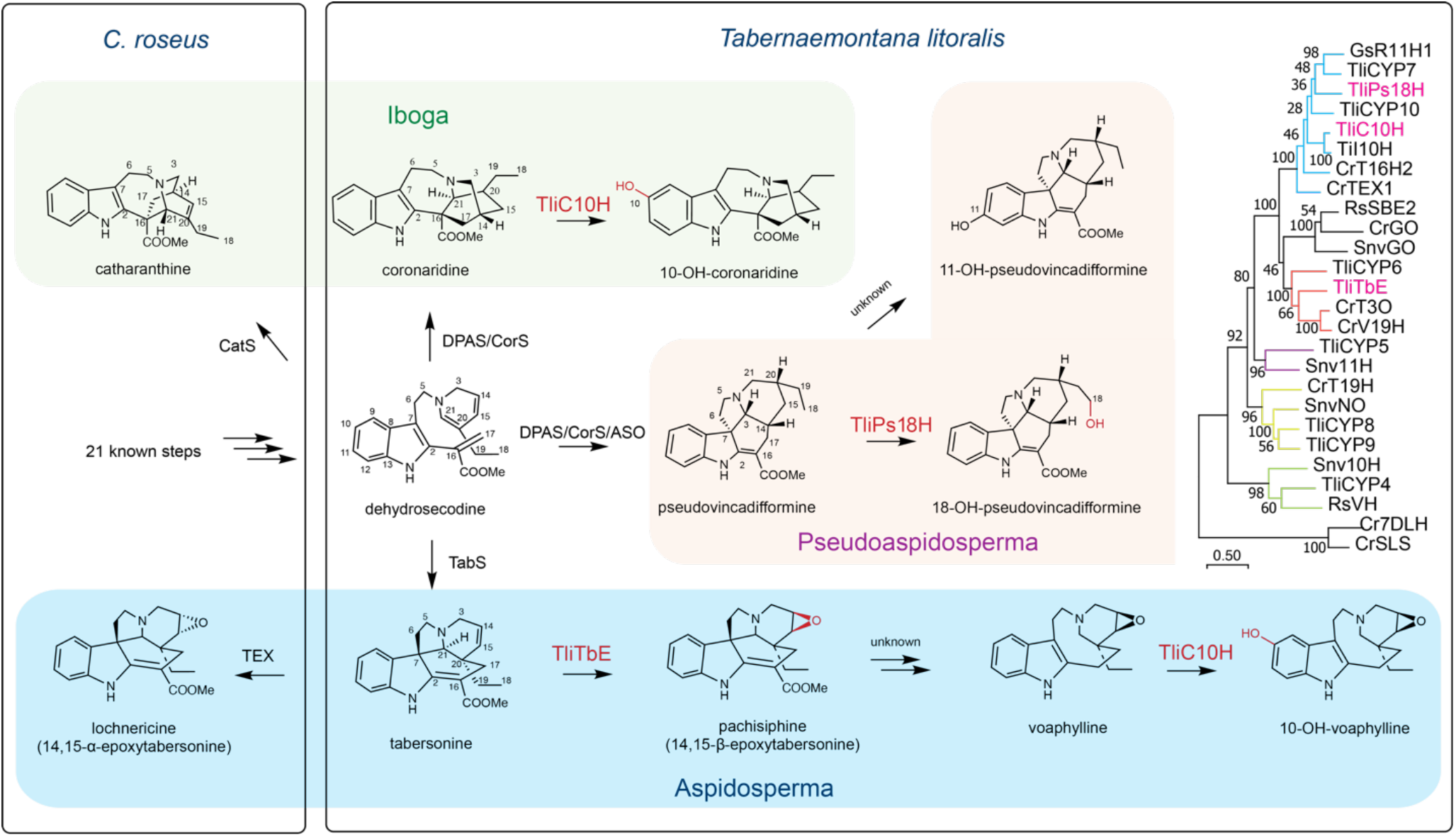
The biosynthetic pathways and CYP phylogeny for iboga, aspidosperma, pseudoaspidosperma, and quebrachamine type monoterpenoid indole alkaloids (MIAs) in *Catharanthus roseus* and *Tabernaemontana litoralis*. The enzymes discovered and characterized in this study are shown in red. Evolutionary analyses were conducted in MEGA11. The evolutionary history was inferred by using the Maximum Likelihood method and JTT matrix-based model. The tree with the highest log likelihood is shown. The percentage of trees in which the associated taxa clustered together is shown next to the branches (100 bootstrap replicates). Initial tree(s) for the heuristic search were obtained automatically by applying Neighbor-Join and BioNJ algorithms to a matrix of pairwise distances estimated using the JTT model, and then selecting the topology with superior log likelihood value. The tree is drawn to scale, with branch lengths measured in the number of substitutions per site (scale bar). CYP4-10: CYP candidates screened in this study; R11H1: rankinidine 11-hydroxylase 1; Ps18H: pseudovincadifformine 18-hydroxylase; I10H: ibogamine 10-hydroxylase; T16H2, tabersonine 16-hydroxylase 2; TEX1: tabersonine 14,15-α-epoxidase; TbE: tabersonine 14,15-β-epoxidase; SBE2: sarpagan bridge enzyme 2; GO: geissoschizine oxidase; T3O: tabersonine 3-oxidase; V19H: (+)-vincadifformine 19-hydroxylase; Snv10H: strychnine 10-hydroxylase; Snv11H: β-colubrine 11-hydroxylase; NO: norfluorocurarine 18-hydroxylase; T19H: tabersonine 19-hydroxylase; VH: vinorine hydroxylase; 7DLH: 7-deoxyloganic acid hydroxylase; SLS: secologanine synthase. Cr: *Catharanthus roseus*; Gs: *Gelsemium sempervirens*; Snv: *Strychnos nux-vomica*; Rs: *Rauvolfia serpentina*; Tli: *Tabernaemontana litoralis*; Ti: *Tabernanthe iboga*. The alignment for constructing the phylogenetic tree is included in Supplementary data 1.

Unlike the omnipresent iboga and aspidosperma MIAs in the Apocynaceae family, the pseudoaspidosperma alkaloids are scarce, despite that all three originate from dehydrosecodine intermediate. Due to this reason, little is known about the medicinal value of the pseudoaspidosperma MIAs. In the model plant *Catharanthus roseus* (Madagascar periwinkle) for MIA research, two homologous enzymes hydrolase 1/catharanthine synthase (HL1/CatS) and hydrolase 2/tabersonine synthase (HL2) directly cyclize dehydrosecodine to catharanthine (iboga) and tabersonine (aspidosperma), respectively (Qu, Easson, *et al*., 2018; Caputi *et al*., 2018). Recently, the delicate conversion of dehydrosecodine to pseudovincadifformine, the principal pseudoaspidosperma skeleton, was reported (Kamileen *et al*., 2022). The biosynthesis involves an initial 4+2 cycloaddition by a *Tabernanthe iboga* hydrolase coronaridine synthase (CorS), followed by repetitive oxidoreduction by dihydroprecondylocarpine synthase (DPAS) and an upstream flavoprotein *O*-acetylstemmadenine oxidase (ASO). Interestingly, these enzymes can also produce a pseudovincadifformine isomer coronaridine (iboga) at pH 9.5, whereas at pH 7.5 pseudovincadifformine was the dominate product. *T. iboga* is well known for accumulating iboga alkaloids such as coronaridine, voacangine, ibogamine, and particularly, the opioid ibogaine that derive from coronaridine. Pseudoaspidosperma MIAs however have not been isolated from this plant. It is possible that other mechanism is involved in determining the product outcome or other enzymes are responsible for the observed results.

Previously, we reported the isolation of a new MIA, 18-hydroxypseudovincadifformine from the fruit of *Tabernaemontana litoralis* (milky way tree), together with iboga MIA coronaridine, 19*S*-hydroxycoronaridine, and 3,19-oxidocoronaridine (Qu *et al*., 2016). 18-hydroxypseudovincadifformine was a major alkaloid in *T. litoralis* fruit, indicating that the species encodes enzymes responsible for the biosynthesis of the rare pseudoaspidosperma type MIAs. Further research in this species will facilitate understanding the mechanism controlling the product formation among iboga, aspidosperma and pseudoaspidosperma skeletons.

In this study, we sequence the *T. litoralis* leaf and root transcriptomes to further understand the biosynthesis of pseudoaspidosperma and other MIAs in this plant. Metabolic profiling in leaf and root tissues identify a new MIA 11-hydroxypseudovincadifformine, along with pseudovincadifformine, 18-hydroxypseudovincadifformine, and voaphylline. We further identify and biochemically characterize three CYPs: pseudovincadifformine 18-hydroxylase (TliPs18H), coronaridine/voaphylline 10-hydroxylase (TliC10H), and tabersonine 14,15-β-epoxidase (TliTbE) for the biosynthesis of pachysiphine and voaphylline. With homology modeling and Michaelis-Menten kinetics, we further discuss their structural features responsible for substrate preference and catalysis.

## Results

### The pseudoaspidosperma-rich *T. litoralis* leaf accumulates a new alkaloid 11-hydroxypseudovincadifformine

Compared to the sarpagan, aspidosperma, and iboga alkaloids found across *Tabernaemontana* species and many others in the Apocynaceae family, the documentations of pseudoaspidosperma MIAs are scarce. Previously we studied *T. litoralis* fruit MIA profile, and showed the presence of iboga, aspidosperma, and pseudoaspidosperma MIAs (Qu *et al*., 2016). To further understand its MIA metabolism, we examined the MIA profiles in glasshouse-grown *T. litoralis* leaf and root tissues by liquid chromatography tandem mass spectrometry (LC-MS/MS) and Nuclear Magnetic Resonance (NMR).

*T. litoralis* was prolific in pseudoaspidosperma alkaloids. In addition to 18-hydroxypseudovincadifformine, we further identified the parental alkaloid pseudovincadifformine, a new MIA 11-hydroxypseudovincadifformine, and a known MIA voaphylline as major MIAs (Fig. 2a). The pseudovincadifformine C11-hydroxylation was indicated by the strong deshielding of C11 resonance (δ 152.8) in ^13^C NMR spectra and the Heteronuclear Multiple Bond Correlation (HMBC) cross peaks among the indole atoms (Figure 2a, Table1, and Supplementary figures 1-5). The NMR chemical shifts of pseudovincadifformine, 11-hydroxyvincadifformine, and 18-hydroxyvincadifformine greatly resembled each other, further supporting the identification (Table 1 and 2, Supplementary figures 1-10). Voaphylline NMR spectra agreed with literature, and the β-epoxide configuration was evident from the Nuclear Overhauser Effect Spectroscopy (NOESY) correlations among H14, 15, 17, 18 and 19 (Figure 2a, Table1 and 2, and Supplementary figure 11-16).

**Table 1.** ^13^C NMR chemical shifts of alkaloids in this study.

|  | Pseudo-<br>vincadifformine | 11-OH-Pseudo-<br>vincadifformine | 18-OH-Pseudo-<br>vincadifformine | Voaphylline |  |  | Pachysiphipine (14,15-β-epoxytabersonine) |  | Lochnericine (14,15-α-epoxytabersonine) |
| --- | --- | --- | --- | --- | --- | --- | --- | --- | --- |
|  | This study<br>acetone-*d*<sub>6</sub> | This study<br>acetone-*d*<sub>6</sub> | (Qu *et al.*,<br>2016)<br>acetone-*d*<sub>6</sub> | This study<br>acetone-*d*<sub>6</sub> | This study<br>CDCl<sub>3</sub> | (Rahman *et al.*, 1983)<br>CDCl<sub>3</sub> | This<br>study<br>CDCl<sub>3</sub> | (Walia *et al.*, 2020)<br>CDCl<sub>3</sub> | (Carqueijeiro *et al.*,<br>2018)<br>CDCl<sub>3</sub> |
| 2 | 165.0 | 167.0 | 165.2 | 140.6 | 139.47 | 139.33 | 165.40 | 165.4 | 167.7 |
| 3 | 66.2 | 67.0 | 66.5 | 54.7 | 53.95 | 58.53 | 49.82 | 49.9 | 50.2 |
| 5 | 51.0 | 52.0 | 51.0 | 54.5 | 53.65 | 53.51 | 51.31 | 51.3 | 50.7 |
| 6 | 44.6 | 45.1 | 44.7 | 26.7 | 26.17 | 26.02 | 44.24 | 44.3 | 44.8 |
| 7 | 55.4 | 56.6 | 55.3 | 109.5 | 109.87 | 109.68 | 54.79 | 54.8 | 54.9 |
| 8 | 137.9 | 137.5 | 137.8 | 129.4 | 128.66 | 128.55 | 137.88 | 137.5 | 137.6 |
| 9 | 121.7 | 110.8 | 121.6 | 118.1 | 117.84 | 117.68 | 121.63 | 121.6 | 121.4 |
| 10 | 120.3 | 114.3 | 120.3 | 119.0 | 118.90 | 118.74 | 120.69 | 120.7 | 120.7 |
| 11 | 127.7 | 152.8 | 127.5 | 120.9 | 120.75 | 120.58 | 127.99 | 128.0 | 127.8 |
| 12 | 109.6 | 110.8 | 109.4 | 111.0 | 110.17 | 110.05 | 109.51 | 109.5 | 109.4 |
| 13 | 144.0 | 140.1 | 144.1 | 137.0 | 135.68 | 135.59 | 143.27 | 143.3 | 143.0 |
| 14 | 36.6 | 37.5 | 36.8 | 52.5 | 52.44 | 52.30 | 52.31 | 52.3 | 54.0 |
| 15 | 32.6 | 33.3 | 33.0 | 59.6 | 59.47 | 59.34 | 56.49 | 56.5 | 57.3 |
| 16 | 95.7 | 95.5 | 95.9 | 23.7 | 23.39 | 23.25 | 91.39 | 91.4 | 90.7 |
| 17 | 26.6 | 27.5 | 26.4 | 37.0 | 36.65 | 32.32 | 23.60 | 23.5 | 23.3 |
| 18 | 11.8 | 12.7 | 60.1 | 7.6 | 7.53 | 7.37 | 7.36 | 7.4 | 7.24 |
| 19 | 28.7 | 29.4 | 39.4 | 33.0 | 32.45 | 36.52 | 26.65 | 26.6 | 24.4 |
| 20 | 35.7 | 36.5 | 30.3 | 34.4 | 33.76 | 33.64 | 37.35 | 37.4 | 41.1 |
| 21 | 54.5 | 55.4 | 54.9 | 59.1 | 58.74 | 53.80 | 71.32 | 71.4 | 67.6 |
| C=O | 167.6 | 168.5 | 167.6 | - | - | - | 169.11 | 169.1 | 168.9 |
| OCH<sub>3</sub> | 50.1 | 50.8 | 50.2 | - | - | - | 51.18 | 51.2 | 51.1 |

**Table 2.**
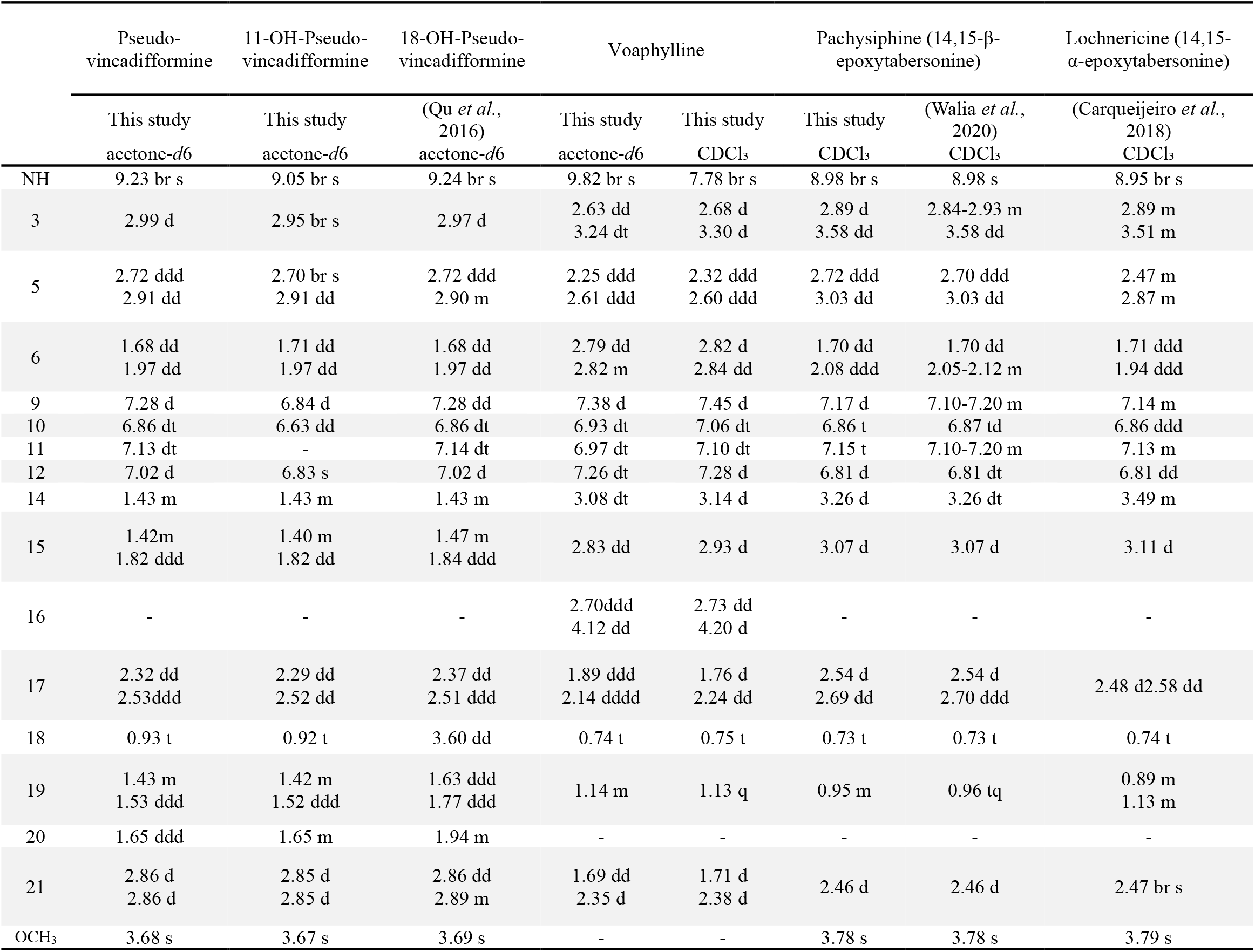
^1^H NMR chemical shifts of alkaloids in this study.

**Figure 2.**
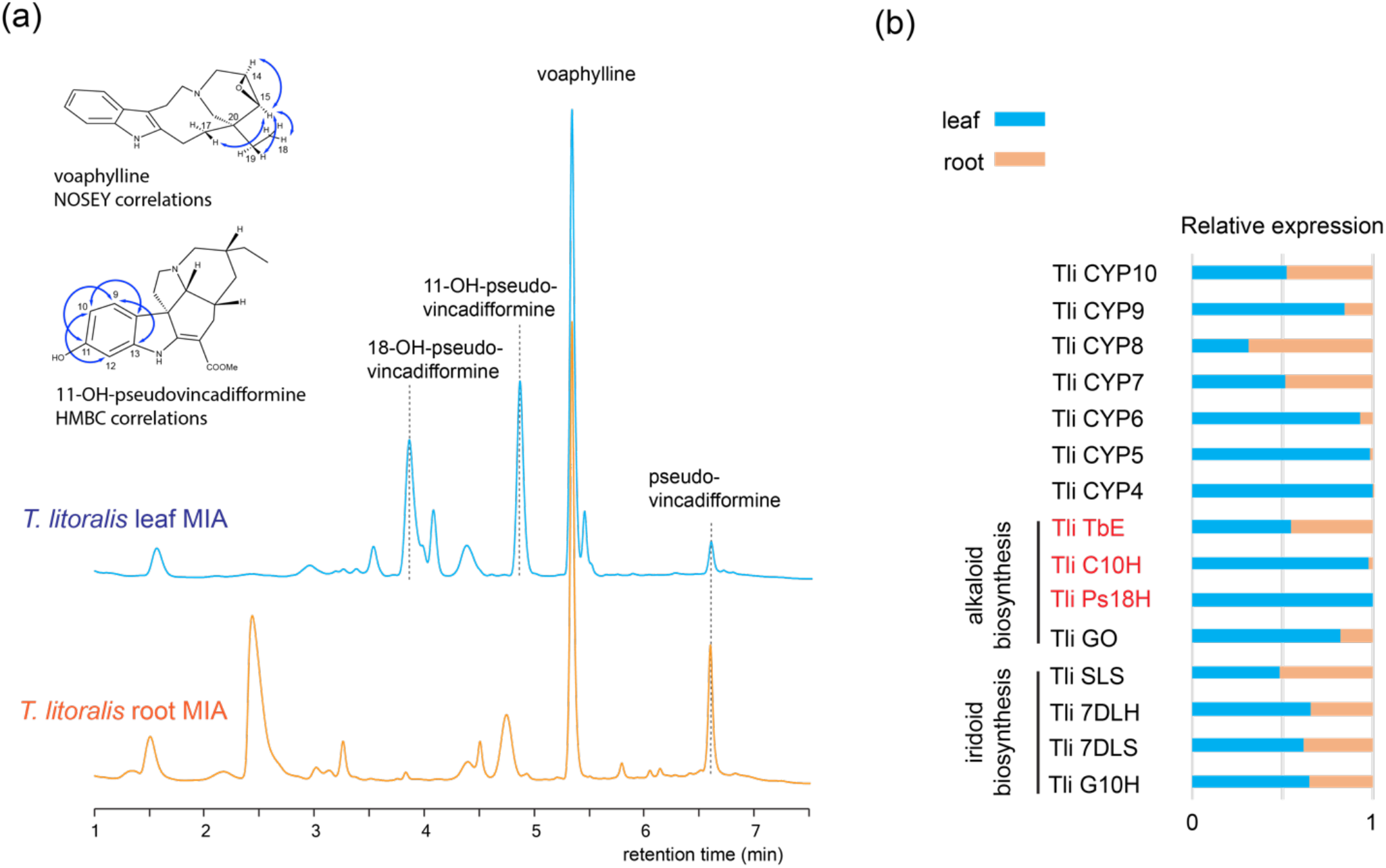
*Tabernaemontana litoralis* leaf and root alkaloid profiles (a) and the relative expression of CYP candidates in leaf and root. (**a**) Liquid Chromatography Tandem Mass Spectrometry (LC-MS/MS) at UV 280 nm indicated differences in alkaloid accumulation in leaf and root, including a new alkaloid 11-hydroxypseudovincadifformine. The alkaloids were identified by standards and Nuclear Magnetic Resonance (NMR) experiments from purified compounds. The Heteronuclear Multiple Bond Correlation (HMBC) cross peaks (blue arrows) were consistent with pseudovincadifformine 11-hydroxylation. The Nuclear Overhauser Effect Spectroscopy (NOESY) experiment (blue arrows) indicated β-epoxide stereochemistry in voaphyllione. The NMR experiments are found in Table 1 and 2, and Supplementary figures 1-22. While voaphylline was present in both leaf and root, 11- and 18-hydroxypseudovincadifformine only accumulated in leaf. (**b**) Relative expressions of the ten CYP candidates and other MIA biosynthetic CYPs in leaf and root. The relative expression was calculated based on their Transcripts Per Million reads (TPM) from Illumina sequencing of respective tissues. The expression data is found in Supplementary table 1.

Compared to the fruit tissue and many other *Tabernaemontana* spp. (Gonçalves *et al*., 2024; Farzana *et al*., 2024), *T. litoralis* leaf and root lacked the sarpagan and iboga alkaloids. Instead, their biosynthetic flux mainly flowed to the pseudoaspidosperma and aspidosperma skeletons. While pseudovincadifformine was hydroxylated at C11 and C18 in leaf, these two alkaloids were not detected in root, where pseudovincadifformine remained unmodified (Fig. 2a). In both tissues, voaphylline (quebrachamine type) was the most abundant MIA. This MIA derives from tabersonine (aspidosperma) through unknown enzymatic steps involving 14,15-epoxidation, carbomethoxyl elimination and perhaps a concomitant C7-C21 fission (Fig. 1).

### *T. litoralis* leaf and root transcriptomes reveal conserved biosynthesis for aspidosperma and pseudoaspidosperma alkaloids

We sequenced the *T. litoralis* leaf and root transcriptomes to further understand MIA biosynthesis in this plant. Sequence alignments with known MIA biosynthetic genes from *C. roseus* identified all biosynthetic genes for forming dehydrosecodine, iboga (coronaridine), aspidosperma (tabersonine), and pseudoaspidosperma (pseudovincadifformine) alkaloids (Fig.1 and Supplementary table 1). Although *T. litoralis* and *C. roseus* belong to two separate Apocynaceae tribes Tabernaemontaneae and Vinceae, respectively, their iridoid and early MIA biosynthetic enzymes were highly conserved, sharing 84-95 % amino acid identity (Supplementary table 1). Less conserved enzymes (63-73% amino acid identity) included strictosidine synthase (STR), strictosidine β-glucosidase (SGD), stemmadenine *O*-acetyltransferase (SAT), *O*-acetylstemmadenine oxidase (ASO), and dihydroprecondylocarpine synthase (DPAS) (Supplementary table 1). In *C. roseus*, dehydrosecodine is directly cyclized to catharanthine and tabersonine (Fig.1), and the latter is further transformed to vindoline. With CorS, ASO, and DPAS, both coronaridine and pseudovincadifformine may form at neutral or basic pH (Fig. 1). We identified both TabS and CorS orthologs in *T. litoralis* leaf and root transcriptomes, responsible for tabersonine and pseudovincadifforming biosynthesis. These two hydrolases shared 84% and 95% amino acid identity with CrTabS and TiCorS, respectively. *T. litoralis* leaf and root accumulated pseudoaspidosperma but iboga MIAs, while iboga MIAs were major MIAs in fruit (Qu *et al*., 2016). Together with the absence of pseudoaspidosperma alkaloids in *T. iboga*, the results suggest that other mechanism might be involved for determining the ratio of iboga/pseudoaspidosperma products from the same dehydrosecodine intermediate.

### *In vivo* screening identifies three *T. litoralis* CYPs for iboga, aspidosperma, pseudoaspidosperma, and quebrachamine MIA oxygenation

In leaf, root, and fruit tissues of *T. litoralis*, the oxygenated MIAs such as voaphylline, 11/18-hydroxypseudovincadifformine, and 19-hydroxycoronaridine (hayneanine) accumulated. CYPs are well-known for compound oxygenation. Therefore, we searched the transcriptomes using known MIA modifying CYPs, and selected the 10 most expressed CYPs in *T. litoralis* leaf (Supplementary table 1). The phylogenetic analysis (Fig. 1) indicated their close evolutionary relationships with MIA indole hydroxylases, such as tabersonine 16-hydroxylase (CrT16H), and side-chain hydroxylases, such as norfluorocurarine 18-hydroxylase (SnvNO) and vincadifformine 19-hydroxylase (V19H) (Hong *et al*., 2022; Williams *et al*., 2022; Williams *et al*., 2019). While some CYPs’s expressions were leaf dominant, other CYP candidates showed balanced expression in both leaf and root. The four CYPs for the formation of MIA precursor secologanin: geraniol 8-hydroxylase (G8H), 7-deoxyloganetic acid synthase (7DLS), 7-deoxyloganic acid hydroxylase (7DLH), and secologanine synthase (SLS) were well expressed in both leaf and root (Fig. 2b).

We then screened baker’s yeast expressing these 10 CYPs by feeding the cultures with 28 MIA substrates encompassing a variety of skeletons (Supplementary table 2). Our screening successfully identified the highest expressed candidates CYP1-3 as the pseudovincadifformine 18-hydroxylase (TliPs18H), coronaridine/voaphylline 10-hydroxylase (TliC10H), and tabersonine 14,15-β-epoxidase (TliTbE) with Genbank accessions PP993475-993477. The remaining CYP4-10 did not show activity with tested substrates.

### The tabersonine 14,15-β-epoxidase (TliTbE) is responsible for pachysiphine and voaphylline biosynthesis

In *C. roseus*, two homologous CYPs, CrTEX1 and CrTEX2, are responsible for the 14,15-α-epoxidation of tabersonine in root and arial tissues, resulting in the formation of lochnericine (14,15-α-epoxitabersonine, Fig. 1) (Carqueijeiro *et al*., 2018). The oxidation product generated by TliTbE with tabersonine displayed a similar MS/MS fragmentation pattern to lochnericine but exhibited a different LC retention time (Fig. 3a and b). After purifying the product from yeast fermentation broth, we confirmed that the epoxidation by TliTbE occurs in the opposite configuration to lochnericine, resulting in the formation of pachysiphine (14,15-β-epoxitabersonine, Fig. 1). The C14,15 alkene resonance signals were replaced by those consistent with an epoxide (δ 52.3/56.5, Table 1, Supplementary figure 17-21.). A NOESY experiment clearly revealed H14,15 cross peaks with H18, H19, H21, and H3α hydrogens, indicating the β-epoxide configuration (Figure 3b, Supplementary figure 21). Pachysiphine is not a common MIA. However, the elimination of the carbomethoxyl group in pachysiphine by an as-yet-unknown process leads to C7-21 fission, forming voaphylline, which is commonly found in *Tabernaemontana* species and many other MIA-producing plants. The expression levels for TliTbE were comparable in both leaf and root (Fig. 2b and Supplementary table 1), consistent with the accumulation of voaphylline as a major MIA in both tissues.

**Figure 3.**
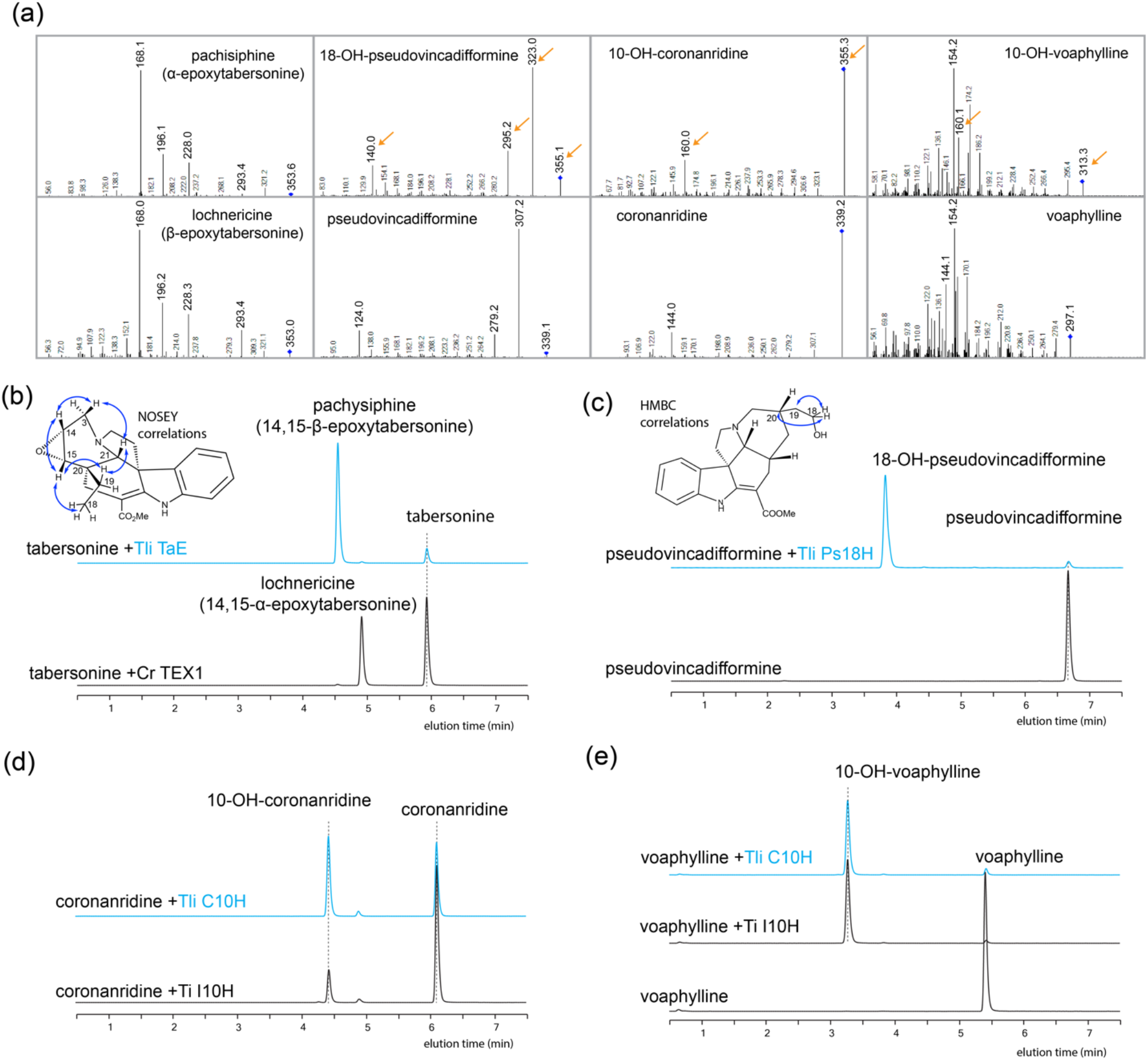
Liquid Chromatography Tandem Mass Spectrometry (LC-MS/MS) experiments demonstrating oxygenation product formations of TliTbE, TliPs18H, and TliC10H. (**a**) MS/MS fragmentations of alkaloids substrates and products. The orange arrows illustrate the increase of 16 amu in parental and daughter ions by oxygenation. (**b**) Yeast expressing TliTbE oxidized tabersonine to pachysiphine (14,15-β-epoxytabersonine) eluted differently from lochnericine (14,15-α-epoxytabersonine) produced by CrTEX1. NOSEY correlations (blue arrows) indicated β-epoxide stereochemistry. (**c**) Yeast expressing TliPs18H hydroxylated pseudovincadifformine at C18. HMBC correlations (blue arrows) indicated cross peaks between H18 hydroxymethylene and H19/H20. (**d**) and (**e**) coronaridine and voaphylline were both hydroxylated at C10 by yeast expressing TliC10H or TiI10H. All chromatograms show Multiple Reaction Monitoring (MRM) traces using these parameters: tabersonine, 337>168; epoxytabersonine 353>168; pseudovincadifformine, 339>307; 18-hydroxypseudovincadifformine, 355>323; coronaridine, 339>144, 10-hydroxycoronaridine, 355>160; voaphylline, 297>154; 10-hydroxyvoaphylline, 313>154.

The Michaelis-Menten kinetics of TliTbE demonstrated a high affinity for tabersonine, as indicated by a significantly lower *K*_*M*_ of 0.277±0.036 μM (Supplementary figure 22), compared to the *K*_*M*_ of 2.08±0.22µM for CrTEX1 (Carqueijeiro *et al*., 2018). To further explore the catalytic mechanisms, we constructed homology models for both epoxidases with AlphaFold 3 (Jumper *et al*., 2021) and studied substrate docking with the Molecular Operating Environment (MOE). In a typical CYP oxidation reaction, molecular oxygen first binds to the heme iron. One oxygen atom is reduced to water, while the other forms a highly reactive ferryl-oxo (Fe^IV^=O) complex (compound I), responsible for CYP’s oxidation activities. In the homology models of TliTbE and CrTEX1, several conserved Thr, Arg, Pro, and Gly residues coordinate heme through hydrogen bonds and van der Waals interactions (Fig. 4). Tabersonine, coordinated mainly by van der Waals interactions, rather adopted opposite docking poses with the ferryl-oxo centers in these two enzymes. This difference in binding orientation leads to the formation of β-epoxytabersonine in TliTbE and α-epoxitabersonine in CrTEX1 (Fig. 4).

**Figure 4.**
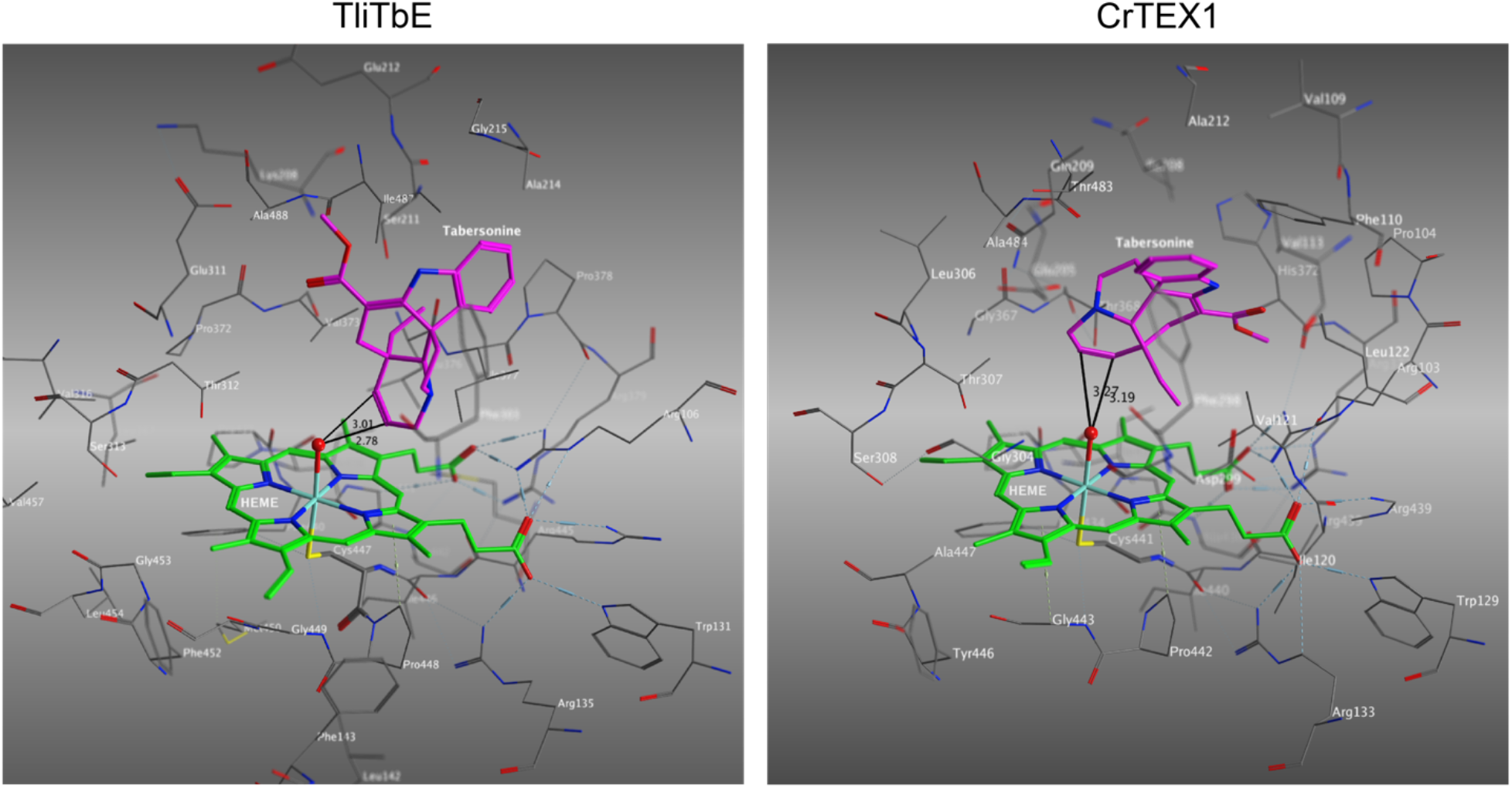
Homology modeling of the active sites of TliTbE and CrTEX1 illustrate the formation of β and α epoxides of tabersonine, respectively. The heme binding residues were conserved between the two CYPs, while the tabersonine binding residues differed significantly. Enzyme models were generated with AlphaFold 3, MOE version 2022.02 was used for substrate docking. Heme is shown in green, and tabersonine is shown in magenta. The distances between the ferryl-oxo oxygen and tabersonine 14,15-double bond are illustrated.

### The pseudovincadifformine 18-hydroxylase (TliPs18H) hydroxylates the ethyl sidechain

The most abundant CYP candidate TliPs18H oxidized pseudovincadifformine, and we identified the product as18-hydroxypseudovincadifformine with a standard previously purified from the fruit. The TliPs18H product showed an increase of 16 amu from the substrate pseudovincadifformine, as well as identical retention time and MS/MS fragments with the standard (Fig. 2a and c). Compared to the root and leaf expressed TliTbE, TliPs18H only expressed in leaf with no detectable transcripts in root (Fig. 2b and Supplementary table 1). The results were consistent with the absence of 18-hydroxypseudovincadifformine in root. Homology modeling for TliPs18H revealed conserved heme binding sites and hydrophobic pocket for substrate binding. Substrate docking with MOE consistently produced top binding poses conducive for closed ferryl-oxo contact with pseudovincadifformine H18. The van der Waals interactions of aliphatic residues oriented pseudovincadifformine for H18 hydroxylation, with an excellent distance of 2.65 Å and 115.0° dihedral angle with the ferrl-oxo center (Fig. 5).

**Figure 5.**
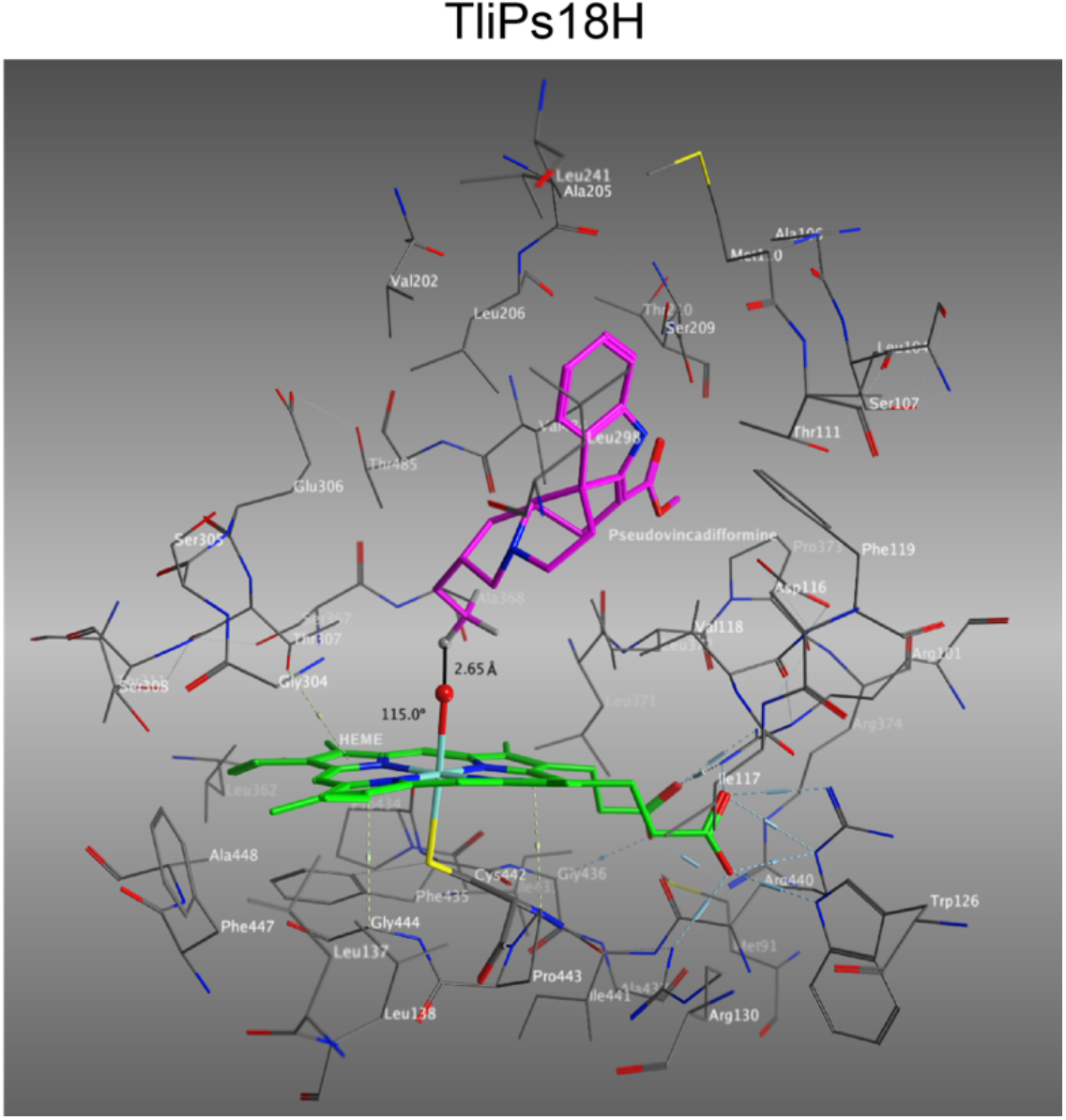
Homology modeling of TliPs18H active site for pseudovincadifformine C18-hydroxylation. The van der Waals interactions of pseudovincadifformine-binding residues facilitate optimal ferryl-oxo-H18 orientation. Enzyme models were generated with AlphaFold 3, MOE version 2022.02 was used for substrate docking. Heme is shown in green, and pseudovincadifformine is shown in magenta. The distance and dihedral angle between the ferryl-oxo oxygen and pseudovincadifformine H18 are illustrated.

### The coronaridine 10-hydroxylase (TliC10H) hydroxylates both coronaridine and voaphylline

In our screening experiments, yeast expressing TliC10H converted both coronaridine and voaphylline into their corresponding oxygenated (+16 amu) products (Fig. 3a, d and e). We confirmed coronaridine 10-hydroxylation using a previously identified *Tabernanthe iboga* ibogamine 10-hydroxylase (TiI10H), which also hydroxylates coronaridine at C10 (Farrow *et al*., 2018). TliC10H and TiI10H share 87% amino acid identity, and both CYPs also oxidized voaphylline to an identical product (Fig. 3e). After purification, we confirmed that the product was hydroxylated at C10. The 10-hydroxyvoaphylline showed resonances for only three aromatic hydrogens in ^1^H-NMR spectrum, with a splitting pattern consistent with C10 or C11 hydroxylation (Supplementary figure 23). Careful comparison with the aromatic resonances from 10- and 11-methoxycoronaridine in literature concluded the hydroxylation on C10 (Chaturvedula *et al*., 2003; González *et al*., 2021).

Homology modeling and substrate docking supported these biochemical results (Fig. 6). In addition to coronaridine and voaphylline, we also studied docking of ibogamine (decarbomethoxylcoronaridine) since it was a substrate of the homologous TiI10H. All three substrates exhibited highly similar binding poses in the hydrophobic active site, despite their structural differences. The binding pocket that accommodated coronaridine’s carbomethoxyl group was similarly occupied by the C18-19-20 ethyl sidechain in voaphylline, which was left empty when ibogamine was docked. The lowest energy poses consistently indicated optimal ferryl-oxo displacement near H11, within 3 Å. This positioning, together with the para position of the indole nitrogen, facilitates hydrogen abstraction and subsequent hydroxyl radical rebounding to the indole radical. The docking of ibogamine to TliC10H and its high sequence similarity to TiI10H suggested that ibogamine is also a substrate of TliC10H.

**Figure 6.**
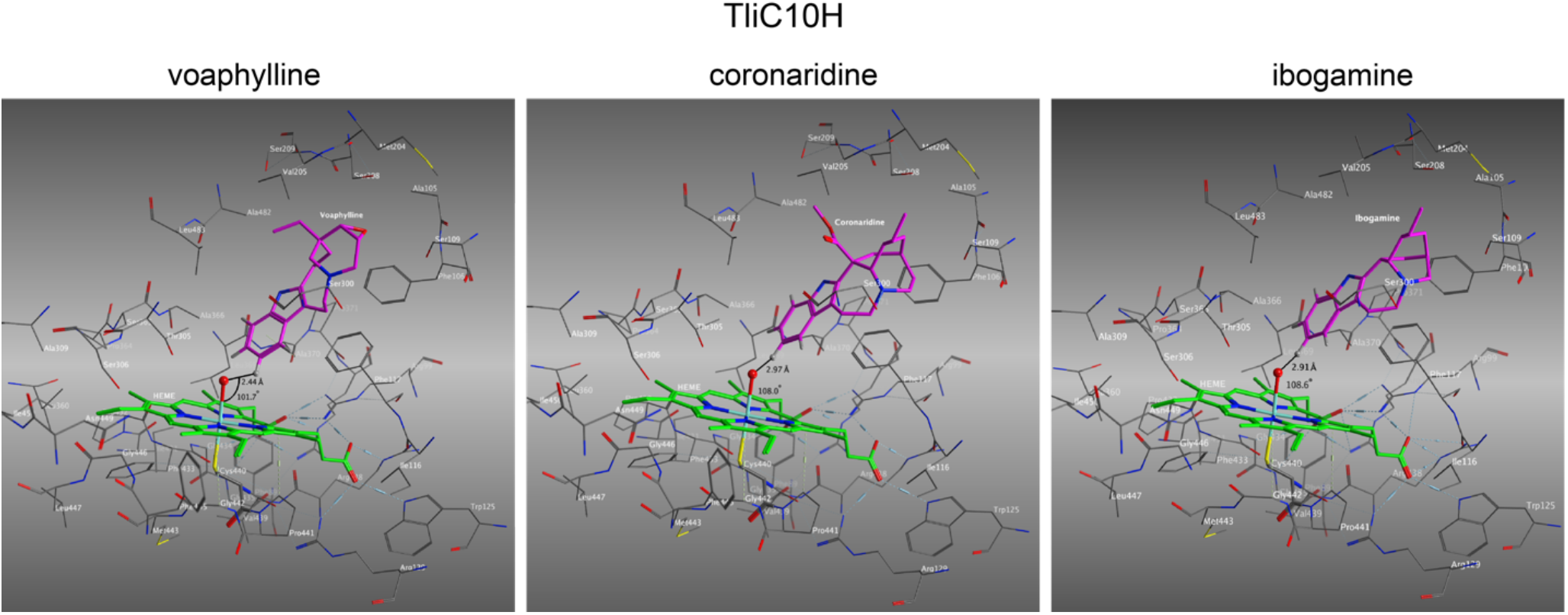
Homology modeling of TliC10H active site for the C10-hydroxylation of coronaridine, voaphylline and ibogamine. All three substrates showed highly similar docking poses for H10 abstraction and subsequent hydroxylation. Enzyme models were generated with AlphaFold 3, MOE version 2022.02 was used for substrate docking. Heme is shown in green, and the substrates are shown in magenta. The distance and dihedral angle between the ferryl-oxo oxygen and the substrate H10 are illustrated.

## Discussion

Extensive studies over the past decade have elucidated the biochemistry and enzymes responsible for a number of medicinal MIAs and their intermediates. Notable examples include the characterization of the over 30-step biosynthetic pathway for the anticancer drug vinblastine from *C. roseus* and the 20-step pathway for the antihypertensive drug ajmaline from *Rauvofia serpentina* (Qu, Safonova, *et al*., 2018; Caputi *et al*., 2018; Guo *et al*., 2024; Dang *et al*., 2018). These discoveries have provided a foundational framework for the biosynthesis of major MIA skeletons, such as corynanthe, sarpagan, strychnos, iboga, aspidosperma, and pseudoaspidosperma, facilitating the study of their more complex and diverse derivatives in other plants. Evolution has shaped specific metabolites in different plant lineages and species, offering fitness advantages in their native environments. The chemotaxonomic distribution of MIAs is evident in families such as Apocynaceae, Rubiaceae, Loganiaceae, and Gelsemiaceae (Martins and Nunez, 2015; Gonçalves *et al*., 2024; Mohammed *et al*., 2021), driven by numerous enzymes evolved uniquely within each species. In *C. roseus*, the most abundant MIAs are catharanthine (iboga type) and vindoline (aspidosperma type), which are not found outside of this small genus. Unique enzymes, such as HL1/CatS and Tabersonine 3-oxygenase (T3O), modify these basic skeletons, diversifying the core MIA metabolism into specific metabolites (Qu *et al*., 2015; Qu, Easson, *et al*., 2018).

In contrast to *C. roseus, T. litoralis* accumulates voaphylline, pseudovincadifformine and its derivatives in both leaf and root. In this work, we identified a new MIA, 11-hydroxypseudovincadifformine. Alongside the previously identified 18-hydroxypseudovincadifformine, which is also unique to this species, we demonstrated that *T. litoralis* is a prolific producer of pseudoaspidosperma MIAs, making it an excellent system for studying the biosynthesis and metabolism of these compounds. While both leaf and root tissues accumulated pseudovincadifformine, yet only in leaf it is further hydroxylated at C11 and C18. Investigating the biological activities of pseudovincadifformine and its hydroxylated derivatives could provide insights into their roles in defense against pathogens and herbivores.

Pseudoaspidosperma MIAs are rare in nature compared to iboga and aspidosperma types. Kamileen et al. demonstrated the biosynthesis of pseudovincadifformine from dehydrosecodine, a highly labile intermediate also involved in the synthesis of iboga and aspidosperma MIAs, via the repeated activities of coronaridine synthase (CorS) and two upstream oxidoreductases from *Tabernanthe iboga* (Kamileen *et al*., 2022). Interestingly, *T. iboga* does not produce pseudovincadifformine but is rich in iboga MIAs such as ibogaine. In this study, we sequenced the leaf and root transcriptomes of *T. litoralis* and identified orthologs of TabS and CorS, as well as other upstream biosynthetic enzymes. Further analysis of these transcriptomes will help understand the enzymes and other factors involved in determining the cyclization outcome of dehydrosecodine. Additionally, we identified and characterized three CYPs for regio- and stereospecific oxidation of iboga, aspidosperma, pseudoaspidosperma, and quebrachamine skeletons, providing new catalysts for accessing various MIA structures.

Homology modeling revealed highly conserved heme-binding sites in all three CYPs in our study (Fig. 4, 5, and 6). However, significant differences lied in substrates binding sites. The hydrophobic binding pocket in TliPs11H oriented the terminal ethyl side chain towards the ferryl-oxo reactive center, despite that TliPs18H shares closer ancestry with MIA indole hydroxylases such as *C. roseus* tabersonine 16-hydroxylase (CrT16H) and *Gelsemium sempervirens* rankinidine 11-hydroxylase (GsR11H) (Fig. 1). The TliPs18H appeared to have evolved specifically in *T. litorali*s. The closest homolog in *C. roseus* genome (Sun *et al*., 2023) and *Rauwolfia tetraphylla* genome (Stander *et al*., 2023) shared 55.0% and 50.3% identity with TliPs11H, respectively. In a closely related species *Voacanga thourasii* (Cuello *et al*., 2022), we could identify a TliPs11H homolog with 78.7% amino acid identity in the genome. None of these species are known to produce pseudoaspidosperma alkaloids, suggesting these homologs may have other physiological roles.

The two tabersonine 14,15-epoxidases CrTEX1 and TliTbE share only 48.2% amino acid identity and belong to two separate clades in our phylogenetic analysis (Fig. 1). As a result, the substrate tabersonine interacts with the heme ferryl-oxo centers of CrTEX1 and TliTbE in opposite orientations, leading to α-epoxidation in CrTEX1 and β-epoxidation in TliTbE. The closer distances between the ferryl-oxo center and the tabersonine 14,15-double bond (2.78/3.01 Å) in TliTbE correlate with its better affinity (*K*_*M*_ 0.277±0.036 µM), compared to CrTEX1 (3.19/3.27 Å) and a reported *K*_*M*_ of 2.08±0.22 µM (Carqueijeiro *et al*., 2018).

After examining other species, we identified two TliTbE orthologs with 93.8% and 94.3% amino acid identity amino acid identity in *Tabernaemontana elegans* and *V. thourasii*, respectively. However, we did not find a homolog with over 55% amino acid identity in the genome of *C. roseus*, nor could we identify a CrTEX1 homolog with over 50% identity in the transcriptomes of *T. litoralis* and *T. elegans*, or the genome of *V. thourasii*. These findings suggest distinct enzyme recruitments between *C. roseus* (Vinceae tribe) and other plants from the Tabernaemontanae tribe of the Apocynaceae family. This distinction is consistent with the presence or absence of specific alkaloids, such as lochnericine and pachysiphine/voaphylline, in these species. The physiological roles of these alkaloids await further investigation.

Interestingly, TliC10H was the second-highest expressed CYP candidate in *T. litoralis* leaf, and its root expression was 2.7% of leaf. However, neither leaf or root contained coronaridine, 10-hydroxycoronaridine, or other iboga MIAs as major alkaloids. Although TliC10H exhibited considerable voaphylline hydroxylation activity, we could not detect 10-hydroxyvoaphylline or putative 10-methoxyvoaphylline in leaf. None of the remaining major leaf alkaloids, including pseudovincadifformine and 11-/18-hydroxypseudovincadifformine, could be attributed to the activity of TliC10H. It is possible that TliC10H has other, as-yet-undetermined activities, or that another factor might be required for the production of coronaridine, providing a suitable substrate for TliC10H under certain conditions. For example, *T. litoralis* fruit accumulated coronaridine, 19*S*-hydroxycoronaridine, 18-hydroxypseudovincadifformine, and tabersonine as major MIAs. The spatial and temporal expression of the biosynthetic enzymes may also contribute to the observed MIA accumulations. The TliC10H ortholog TiI10H from *T. iboga* is known to hydroxylate C10 of ibogamine (decarbomethoxylcoronaridine). While we did not have the ibogamine substrate, it is likely that TliC10H also has ibogamine 10-hydroxylase activity based on substrate docking experiment. Nonetheless, the finding of their voaphylline 10-hydroxylation activity extends our understanding of the substrate preference and product spectrum of these CYPs.

After screening the 10 most expressed CYP candidates in *T. litoralis* leaf, we did not identify the pseudovincadifformine 11-hydroxylase. It is possible that another CYP with lower expression is responsible for the activity, or that this hydroxylation is not catalyzed by a CYP but by another type of oxidase. Further search in the leaf transcriptome may identify this oxidase. Further research is also needed to discover enzymes for pachysiphine decarbomethoxylation and mechanistic insights for dehydrosecodine cyclization, using the *T. litoralis* transcriptomes and those of other MIA-producing plants.

## Materials and Methods

### Plant materials and alkaloid purification

Glasshouse grown *T. litoralis* Fresh leaves (200 g) were submerged in methanol to yield total crude extract, which was evaporated under vacuum and dissolved in 1 M HCl. After extraction with ethyl acetate, the aqueous phase was basified to above pH 7, and extracted by ethyl acetate to yield the total alkaloids. The total alkaloids were separated by thin layer chromatography (Sicica gel60 F254, Millipore Sigma, Rockville, MD, USA) with solvents ethyl acetate and methanol (9:1, v/v), which yielded voaphylline, 11-hydroxypseudovincadifformine, 18-hydroxypseudovincadifformine, and pseudovincadifformine. Other alkaloid standards were obtained from previous research (Qu *et al*., 2016).

### mRNA extraction and transcriptome assembly

*T. litoralis* young leaf and root tissues (100 mg) from glasshouse grown plants were ground into fine powder in liquid nitrogen. The total RNA were extracted using a Mornarch® RNA Cleanup Kit (New England Biolabs, Ipswich, MA, USA) according to the manufacture’s protocol. The RNA was sequenced using Illumina NovaSeq 25M reads at the Atlantic Cancer Research Institute (Moncton, NB, Canada). The transcriptomes were assembled using Trinity scripts and analyzed using CLC Genomic Workbench 20.0.4 (Qiagen, Redwood City, CA, USA).

### LC-MS/MS and NMR

The samples were analyzed by Ultivo Triple Quadrupole LC-MS/MS system from Agilent (Santa Clara, CA, USA), equipped with an Avantor® ACE® UltraCore C18 2.5 Super C18 column (50×3 mm, particle size 2.5 µg,) as well as a photodiode array detector and a mass spectrometer. For alkaloid analysis, the following solvent systems were used: Solvent A, Methanol: Acetonitrile: Ammonium acetate 1 M, water at 29:71:2:398; solvent B, Methanol: Acetonitrile: Ammonium acetate 1 M: water at 130:320:0.25:49.7. The following linear elution gradient was used: 0-5 min 80% A, 20% B; 5-5.8 min 1% A, 99% B; 5.8-8 min 80% A, 20% B; the flow during the analysis was constant and 0.6 ml/min. The photodiode array detector range was 200 to 500 nm. The mass spectrometer was operated with the gas temperature at 300°C and gas flow of 10 L/min. Capillary voltage was 4 kV from m/z 100 to m/z 1000 with scan time 500 ms and the fragmentor performed at 135 V with positive polarity. The MRM mode was operated as same as scan mode with the adjusted precursor and product ion. NMR spectra were recorded on an Agilent 400 MR and a Bruker Avance AV I 600 digital NMR spectrometer in acetone-*d6* or CDCl_3_.

### Cloning

*T. litoralis* CYP candidates were amplified from the total leaf cDNA with primers (Supplementary table 3). TliTbE and TliCYP5 were ligated into pESC-Leu vector with CrCPR previously cloned, within ApaI/SalI sites. The remaining CYPs were ligated into BamHI/SalI sites in the same vector. The clones were mobilized to *Saccharomyces cerevisiae* strain BY4741 (MATα his3Δ1 leu2Δ0 met15Δ0 ura3Δ0 YPL154c::kanMX4) with standard lithium acetate/ polyethylene glycol transformation procedure.

### Yeast biotransformation

Single colonies of the yeasts carrying various CYPs were inoculated in 1 mL synthetic complete (SC)-Leu media with 2% (w/v), and incubated at 30 °C, 200 rpm overnight. The yeasts were pelleted by centrifugation, washed once with water, resuspended in 1 mL SC-Leu media with 2% (w/v) galactose, and incubated at 30 °C, 200 rpm overnight. The yeasts were pelleted by centrifugation, and resuspended in 0.2 mL 20 mM Tris-HCl supplemented with 0.2-2 μg alkaloids. The biotransformation took place at 30 °C, 200 rpm overnight, which was mixed with equal volume of methanol for LC-MS/MS analysis.

### Yeast microsomes preparation and in vitro kinetics assay

Yeast culture (2 mL) carrying CYP of interest was used to inoculate 200 mL SC-Leu media with with 2% (w/v) glucose, which was incubated at 30 °C, 200 rpm overnight. The cells were pelleted by centrifugation, washed once with water, resuspended in 200 mL SC-Leu media with 2% (w/v) galactose, and incubated at 30 °C, 200 rpm overnight. The pelleted cells were suspended in 4 mL ice cold TES buffer (10 mM Tris-HCl pH 8, 1 mM EDTA, 0.6 M sorbitol), and lysed by beating the cells with glassbeads using a Qiagen tissuelyser II (Qiagen, Germantown, MD, USA). The lysate was centrifuged at 16,000 g at 4 °C for 10 min, and the supernatant was further centrifuged at 100,000 g at 4 °C for 1 hr to afford microsomes. The microsomes were resuspended in ice cold TEG buffer (10 mM Tris-HCl pH 8, 1 mM EDTA, 10% (v/v) glycerol) and stored at −80 °C. The kinetics assay (100 μL) included 100 mM potassium phosphate pH 7.0, 1 mM NADPH, 500 μg microsomal proteins, and tabersonine substrate at various concentrations. The kinetics reactions were carried out at 30 °C for 30 min, then was terminated by adding equal volume of methanol for LC-MS analysis. Triplicated experiments were graphed using GraphPad Prism (9.5.0) (Boston, MA, USA).

### Homology modeling and substrate docking studies

All computational experiments and visualization were carried out with MOE version 2022.02 and AlphaFold 3 on local computers or on the Digital Research Alliance of Canada (DRAC, formerly Compute Canada) Advanced Research Computing Network (alliancecan.ca). All molecular mechanics calculations and simulations were conducted using the AMBER14:EHT forcefield modified for ferryl-oxo bond. Homology models were obtained in AlphaFold using the protein amino acid sequence and heme as the selected ligand. The homology models were consequently transferred to MOE and an oxygen atom was then covalently bound to the Fe atom of heme, 180º from the opposing cysteine residue, using MOE’s builder function. The model was then subject to a QuickPrep using default settings. The formal charge of oxygen was then altered to −1, resulting in a Fe-O carbonyl equivalent. The enzymes were then energy minimized using R-Field and Born solvation, respectively, with a dielectric constant of 1 and an exterior dielectric constant of 80. Using the default MOE-Dock settings, the heme molecule, including the bound oxygen species was selected as the general docking region. Ligands were then modeled in MOE and subject to a conformational search, deleting any conformers with more than a 6-kcal difference in energy compared to the lowest energy conformer. The conformational search file of each ligand was then docked into the given protein-heme complex; for each ligand docking experiment, 17,000 docking poses were initially generated via the Triangle Matcher method and scored by the GBVI/WSA dG function. A subset comprising the best docking poses was refined by the induced fit method, where the bound ligands and active site residues were submitted to local geometry optimization and rescored once more by the GBVI/WSA dG scoring function. The top-scoring docking poses that were geometrically conducive to their respective reaction mechanisms were retained for subsequent analyses. Cartesian coordinates for all homology models and their ligand complexes can be found in the Supplementary data 2.

## Acknowledgements

The authors thank the Chemical Computing Group ULC (www.chemcomp.com) for MOE licenses. This research was enabled in part by support provided by ACENET (www.ace-net.ca) and the Digital Research Alliance of Canada (alliancecan.ca).

