## Supplementary information for "Oxidation of Four Monoterpenoid Indole Alkaloid Classes by Three Cytochrome P450 Monooxygenases from *Tabernaemontana litoralis*"

[*]Corresponding authors

Table of Contents


***Supplementary table 1. T. litoralis leaf alkaloid biosynthetic genes and CYP candidates*** **3**

***Supplementary table 2. Alkaloid substrates used in yeast screening experiment*** **4**

***Supplementary table 3. Primers in this study.*** **5**

***Supplementary figure 1. $^{1}$H NMR spectra for 11-hydroxypseudovincadifformine in acetone-d6.*** **6**

***Supplementary figure 2. $^{13}$C NMR spectra for 11-hydroxypseudovincadifformine in acetone-d6.*** **7**

***Supplementary figure 3. HSQC NMR spectra for 11-hydroxypseudovincadifformine in acetone-d6.*** **8**

***Supplementary figure 4. HMBC NMR spectra for 11-hydroxypseudovincadifformine in acetone-d6.*** **9**

***Supplementary figure 5. COSY NMR spectra for 11-hydroxypseudovincadifformine in acetone-d6.*** **10**

***Supplementary figure 6. $^{1}$H NMR spectra for pseudovincadifformine in acetone-d6.*** **11**

***Supplementary figure 7. HSQC NMR spectra for pseudovincadifformine in acetone-d6.*** **12**

***Supplementary figure 8. HMBC NMR spectra for pseudovincadifformine in acetone-d6.*** **13**

***Supplementary figure 9. NOSEY NMR spectra for pseudovincadifformine in acetone-d6.*** **14**

***Supplementary figure 10. COSY NMR spectra for pseudovincadifformine in acetone-d6.*** **15**

***Supplementary figure 11. $^{1}$H NMR spectra for voaphylline acetone-d6.*** **16**

***Supplementary figure 12. $^{13}$C NMR spectra for voaphylline acetone-d6.*** **17**

***Supplementary figure 13. HSQC NMR spectra for voaphylline acetone-d6.*** **18**

***Supplementary figure 14. HMBC NMR spectra for voaphylline.*** **19**

***Supplementary figure 15. NOSEY NMR spectra for voaphylline.*** **20**

***Supplementary figure 16. COSY NMR spectra for voaphylline in acetone-d6.*** **21**

***Supplementary figure 17. $^{1}$H NMR spectra for pachisiphine in $CDCl_3$.*** **22**

***Supplementary figure 18. $^{13}$C NMR spectra for pachisiphine in $CDCl_3$.*** **23**

***Supplementary figure 19. HSQC NMR spectra for pachisiphine in $CDCl_3$.*** **24**

***Supplementary figure 20. HMBC NMR spectra for pachisiphine in $CDCl_3$.*** **25**

***Supplementary figure 21. NOSEY NMR spectra for pachisiphine in $CDCl_3$.*** **26**

***Supplementary figure 22. Saturation kinetics for TliTbE.*** **27**

***Supplementary figure 23. 1H NMR spectra for 10-hydroxyvoaphylline and voaphylline in $CDCl_3$.*** **28**

***Supplementary data 1. Protein alignments used for phylogenetic studies.*** **29**

### Supplementary table 1. *T. litoralis* leaf alkaloid biosynthetic genes and CYP candidates

| Query | gene | Lowest E-value | Accession (E-value) | Greatest identity % | Accession (identity %) | leaf TPM | Root TPM |
|---|---|---|---|---|---|---|---|
| TRINITY_DN765_c0_g1_i1 | TliG10H | 0 | CrG10H | 86.6 | CrG10H | 174.40 | 96.37 |
| TRINITY_DN3884_c0_g1_i4 | Tli8HGO | 0 | Cr8HGO | 91.0 | Cr8HGO | 209.85 | 74.81 |
| TRINITY_DN5687_c0_g1_i3 | Tli7DLS | 0 | Cr7DLS | 94.7 | Cr7DLS | 471.53 | 296.44 |
| TRINITY_DN6961_c0_g1_i1 | Tli7DLGT | 0 | Cr7DLGT | 90.7 | Cr7DLGT | 58.82 | 81.90 |
| TRINITY_DN6632_c0_g1_i2 | Tli7DLH | 0 | Cr7DLH | 90.0 | Cr7DLH | 169.24 | 88.60 |
| TRINITY_DN160_c0_g4_i2 | TliLAMT | 0 | CrLAMT | 84.4 | CrLAMT | 1134.75 | 309.89 |
| TRINITY_DN6588_c0_g1_i1 | TliSLS | 0 | CrSLS | 92.4 | CrSLS | 853.71 | 895.90 |
| TRINITY_DN127_c0_g1_i1 | TliTDC | 0 | CrTDC | 85.8 | CrTDC | 328.23 | 92.40 |
| TRINITY_DN25172_c0_g1_i1 | TliSTR | 1.9417E-175 | CrSTR | 71.8 | CrSTR | 198.03 | 113.76 |
| TRINITY_DN626_c0_g1_i12 | TliSGD | 0 | CrSGD | 67.8 | CrSGD | 102.50 | 34.91 |
| TRINITY_DN293_c0_g1_i2 | TliGS | 0 | CrGS | 87.9 | CrGS | 510.79 | 574.01 |
| TRINITY_DN5537_c0_g1_i2 | TliGO | 0 | CrGO | 91.6 | CrGO | 2094.01 | 471.45 |
| TRINITY_DN86_c0_g3_i1 | TliRedox1 | 1.8285E-178 | CrReDOX1 | 85.3 | CrReDOX1 | 636.96 | 126.06 |
| TRINITY_DN944_c0_g1_i3 | TliRedox2 | 0 | CrReDOX2 | 83.8 | CrReDOX2 | 783.02 | 77.27 |
| TRINITY_DN4819_c0_g1_i3 | TliSAT | 0 | CrSAT | 76.0 | CrSAT | 98.05 | 22.32 |
| TRINITY_DN11_c0_g2_i4 | TliASO1 | 0 | CrASO | 68.8 | CrASO | 110.99 | 43.63 |
| TRINITY_DN11_c0_g2_i1 | TliASO2 | 0 | CrASO | **62.8** | CrASO | 372.35 | 59.65 |
| TRINITY_DN86_c1_g1_i3 | TliDPAS1 | 1.7183E-180 | CrDPAS | 78.4 | CrDPAS | 219.26 | 95.32 |
| TRINITY_DN86_c0_g1_i25 | TliDPAS2 | 1.3569E-163 | CrDPAS | 66.6 | CrDPAS | 47.33 | 301.98 |
| TRINITY_DN1524_c0_g2_i1 | TliHL1/CorS | 0 | CrHL3 | 78.4 | CrHL3 | 942.53 | 170.62 |
| TRINITY_DN1524_c0_g1_i1 | TliHL2/TabS | 0 | CrHL3 | 83.6 | CrHL2 | 104.72 | 639.01 |
| TRINITY_DN1782_c0_g1_i48 | TliPs11H | 9.924E-176 | CrTEX1 | 58.2 | CrT16H | 2225.95 | 0.00 |
| TRINITY_DN43_c1_g3_i1 | TliC10H | 1.6471E-152 | CrT16H | 59.6 | CrT16H | 1810.48 | 49.74 |
| TRINITY_DN3546_c0_g1_i4 | TliTbE | 0 | CrT3O | 55.7 | CrT3O | 307.51 | 251.54 |
| TRINITY_DN4776_c0_g1_i1 | TliCYP4 | 1.5448E-162 | RsVH | 46.8 | RsVH | 174.23 | 0.65 |
| TRINITY_DN878_c0_g1_i1 | TliCYP5 | 5.1687E-109 | CrTEX1 | 53.3 | CrT19H | 124.08 | 2.00 |
| TRINITY_DN3988_c0_g1_i1 | TliCYP6 | 1.3096E-167 | CrT3O | 55.6 | CrBIS1 | 78.82 | 6.49 |
| TRINITY_DN1782_c0_g1_i52 | TliCYP7 | 0 | CrT16H | 57.1 | CrT16H | 79.23 | 75.36 |
| TRINITY_DN1863_c0_g2_i2 | TliCYP8 | 1.8393E-175 | CrT19H | 77.8 | CrNMT | 55.40 | 122.17 |
| TRINITY_DN1863_c0_g1_i1 | TliCYP9 | 0 | CrT19H | 53.0 | CrT19H | 52.33 | 10.12 |
| TRINITY_DN43_c0_g1_i1 | TliCYP10 | 0 | CrTEX1 | 59.4 | CrT16H | 41.74 | 37.45 |

### Supplementary table 2. Alkaloid substrates used in yeast screening experiment

| heteroyohimbine | corynanthe | yohimbe | iboga | Aspidosperma | sarpagan | akuammiline | other |
|---|---|---|---|---|---|---|---|
| Ajmalicine | Corynantheidine | Yohimbine | Catharanthine | Tabersonine | Pericyclivine | Picrinine | Vincamine |
| | hirsuteine | corynanthine | Coronaridine | minovine | pseudoakuammigine | Rhazimal | voaphylline |
| | | Reserpic acid | | Vincaminoreine | perivine | Picralinal | ellipticine |
| | | | | Vincadifformine | Strictamine | | akuammicine |
| | | | | Minovincinine | Akuammidine | | pseudovincadifformine |
| | | | | | vomilenine | | apparicine |

**Supplementary table 3. Primers in this study.**

| # | primer | sequence (5'-3') |
|---|---|---|
| 1 | TliPs18H-BamHI-F | ATAGGATCCATGGAGTTCTTCACTGCCTTATG |
| 2 | TliPs18H-SalI-R | ATAGTCGACGACATGAGAGGAATGACTATAAGGA |
| 3 | TliC10H-BamHI-F | ATAGGATCCATGGAGCTTATCGTTTCCCTCT |
| 4 | TliC10H-SalI-R | ATAGTCGACTTTTTTCATGAATGAACGAGAATAGACC |
| 5 | TliTbE-ApaI-F | AGGGCCCATGGAGCTTCAGAACTTACCCTT |
| 6 | TliTbE-SalI-R | ATAGTCGACGCATGATTTATCAAGAGTCGGAATATCG |
| 7 | TliCYP4-BglII-F | ATAAGATCTATGGATATCCTTTCAAGCATCTTGG |
| 8 | TliCYP4-SalI-R | ATAGTCGACCAGTTTCTGATAGAGCTCTAGTGGAA |
| 9 | TliCYP5-ApaI-F | AGGGCCCATGGGGTTTCAATGGGAAATTGTGCTT |
| 10 | TliCYP5-XhoI-R | AATACTCGAGGTCTTCAGTCATAAACACTTCGTAT |
| 11 | TliCYP6-BamHI-F | ATAGGATCCATGGAGTTGCTTTATCTGTTCGAC |
| 12 | TliCYP6-SalI-R | ATAGTCGACCATGGAAACATCATAGGGTGTGG |
| 13 | TliCYP7-BamHI-F | ATAGGATCCTCAAACCTCATGGAGATCATCTTCC |
| 14 | TliCYP7-SalI-R | ATAGTCGACTTGTAAATAAGAACGAGAGTAAGGAA |
| 15 | TliCYP8-BamHI-F | ATAGGATCCGATACTTTCCAGCAAATGATGTCCA |
| 16 | TliCYP8-SalI-R | ATAGTCGACCATTGAATAAGGAGTGGCAACAAC |
| 17 | TliCYP9-BamHI-F | ATAGGATCCAGAGAGAAGCAAATGGTGTTTCT |
| 18 | TliCYP9-SalI-R | ATAGTCGACATCCACCGATTTTGTTCCTCCAT |
| 19 | TliCYP10-BamHI-F | ATAGGATCCATGGAGGTCGTCTTTCTCATCTGT |
| 20 | TliCYP10-SalI-R | ATAGTCGACTTTTTTCAAAGAAGAAGAGTGATAAAGC |

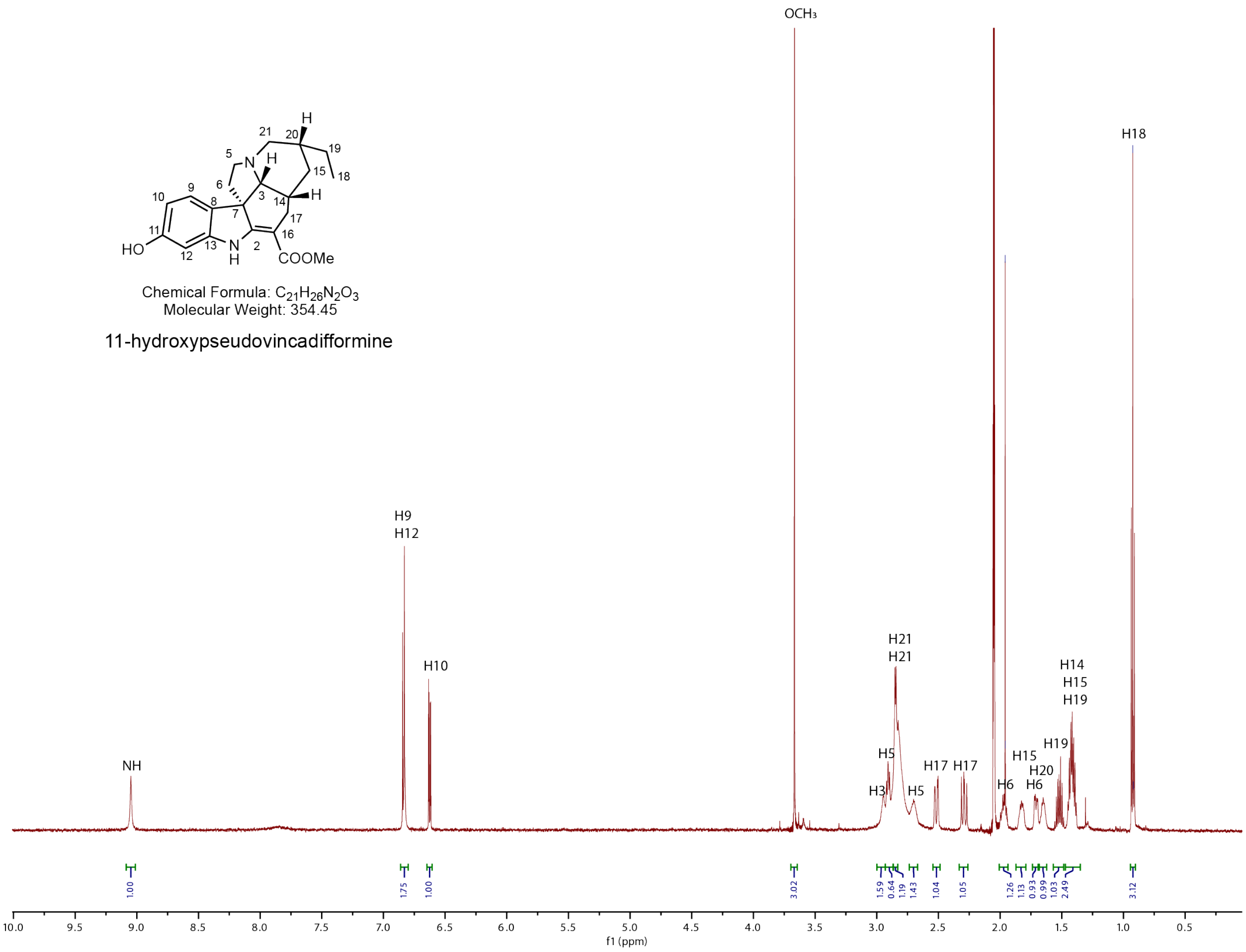


**Supplementary figure 1. $^{1}$H NMR spectra for 11-hydroxypseudovincadifformine in acetone-*d6*.**

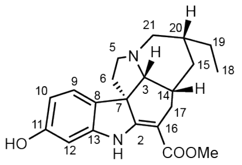


Chemical Formula: $C_{21}H_{26}N_2O_3$
Molecular Weight: 354.45

11-hydroxypseudovincadifformine

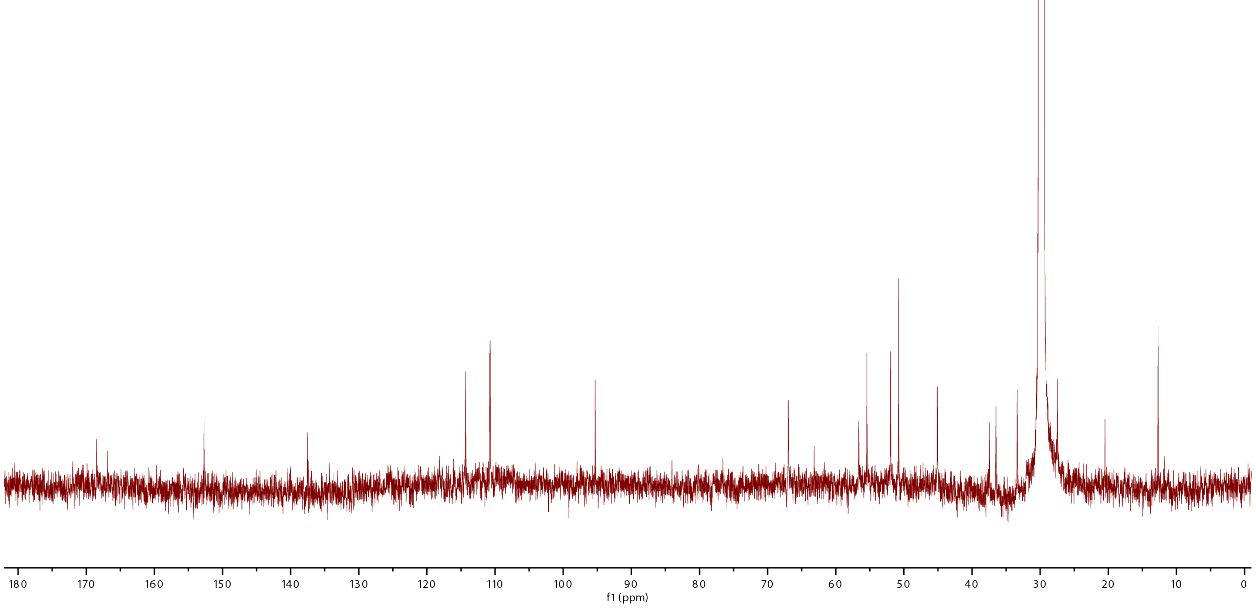


**Supplementary figure 2. $^{13}$C NMR spectra for 11-hydroxypseudovincadifformine in acetone-*d6*.**

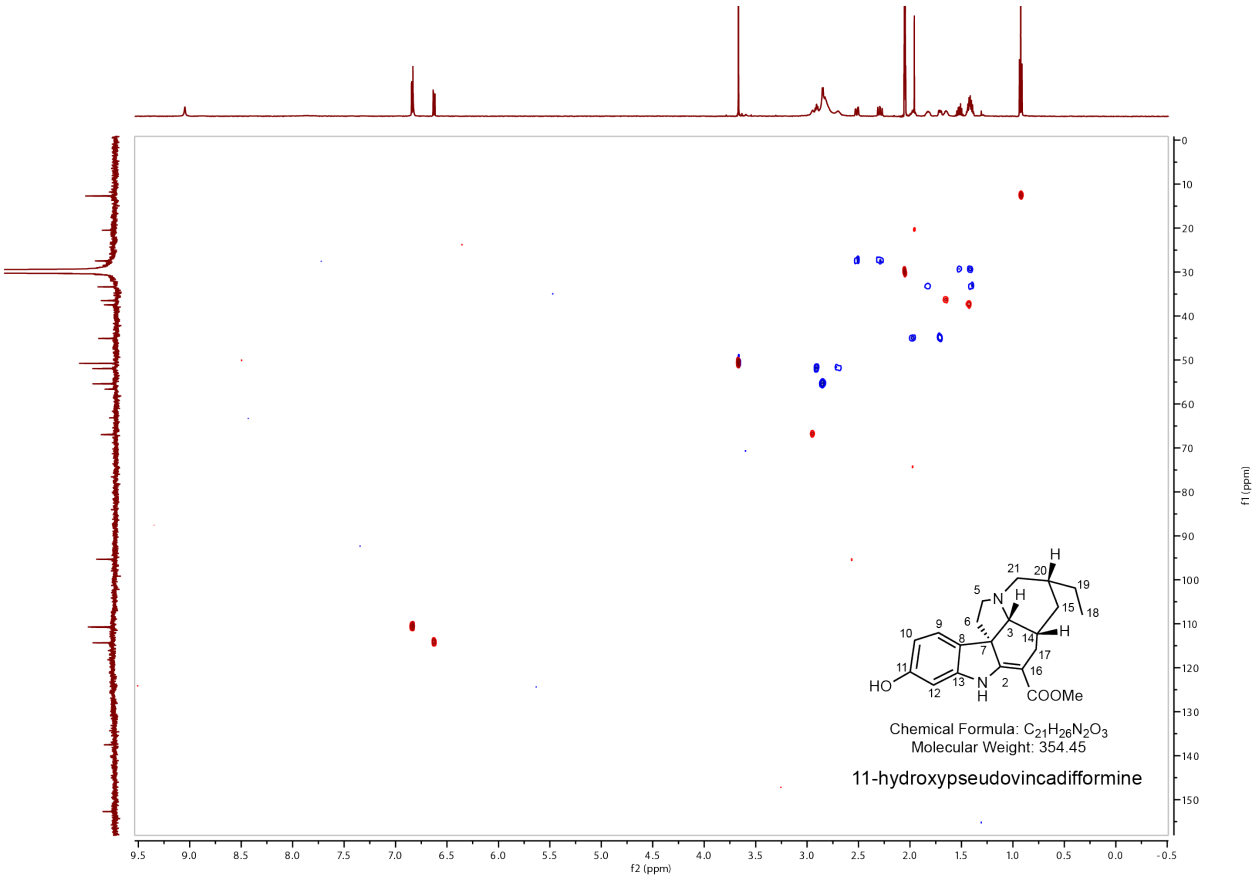


**Supplementary figure 3. HSQC NMR spectra for 11-hydroxypseudovincadifformine in acetone-d6.**

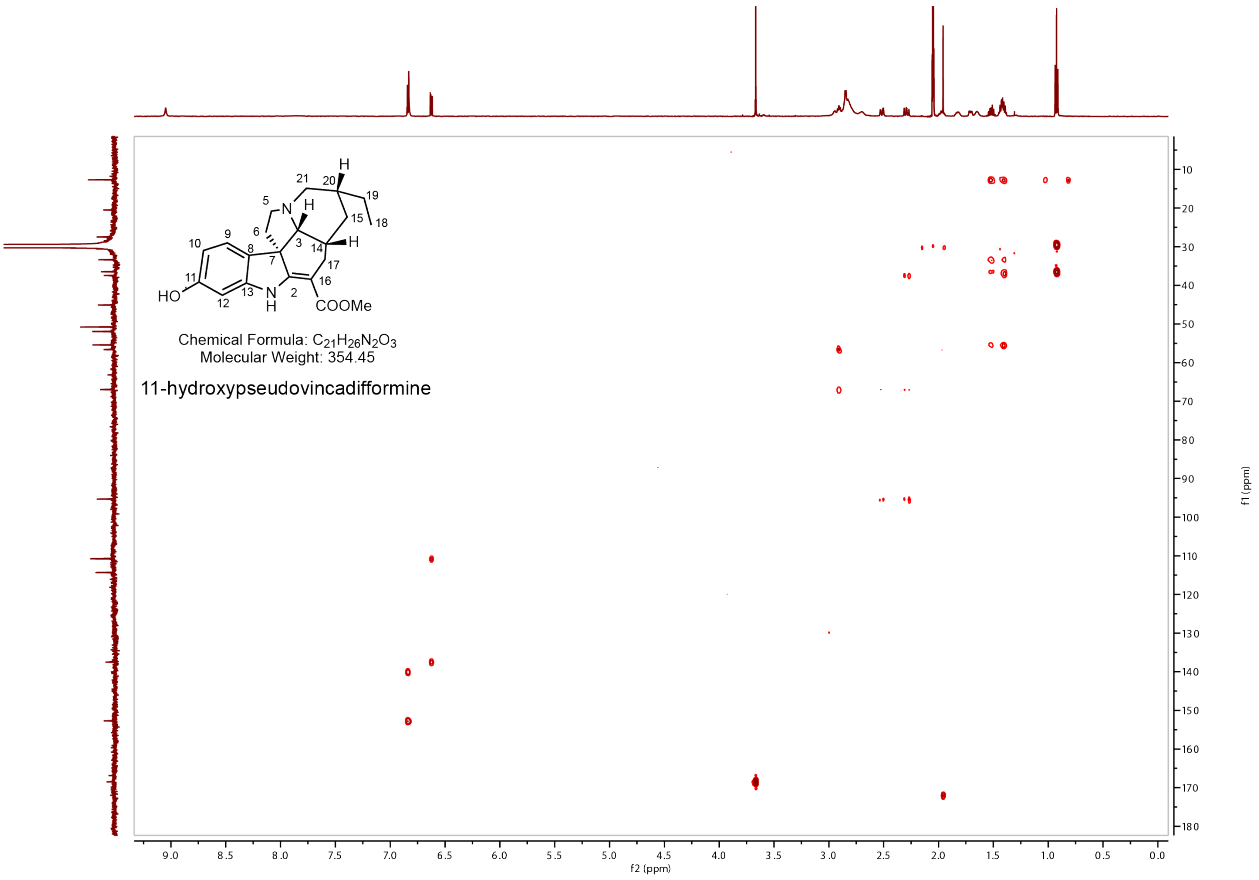


**Supplementary figure 4. HMBC NMR spectra for 11-hydroxypseudovincadifformine in acetone-*d6*.**

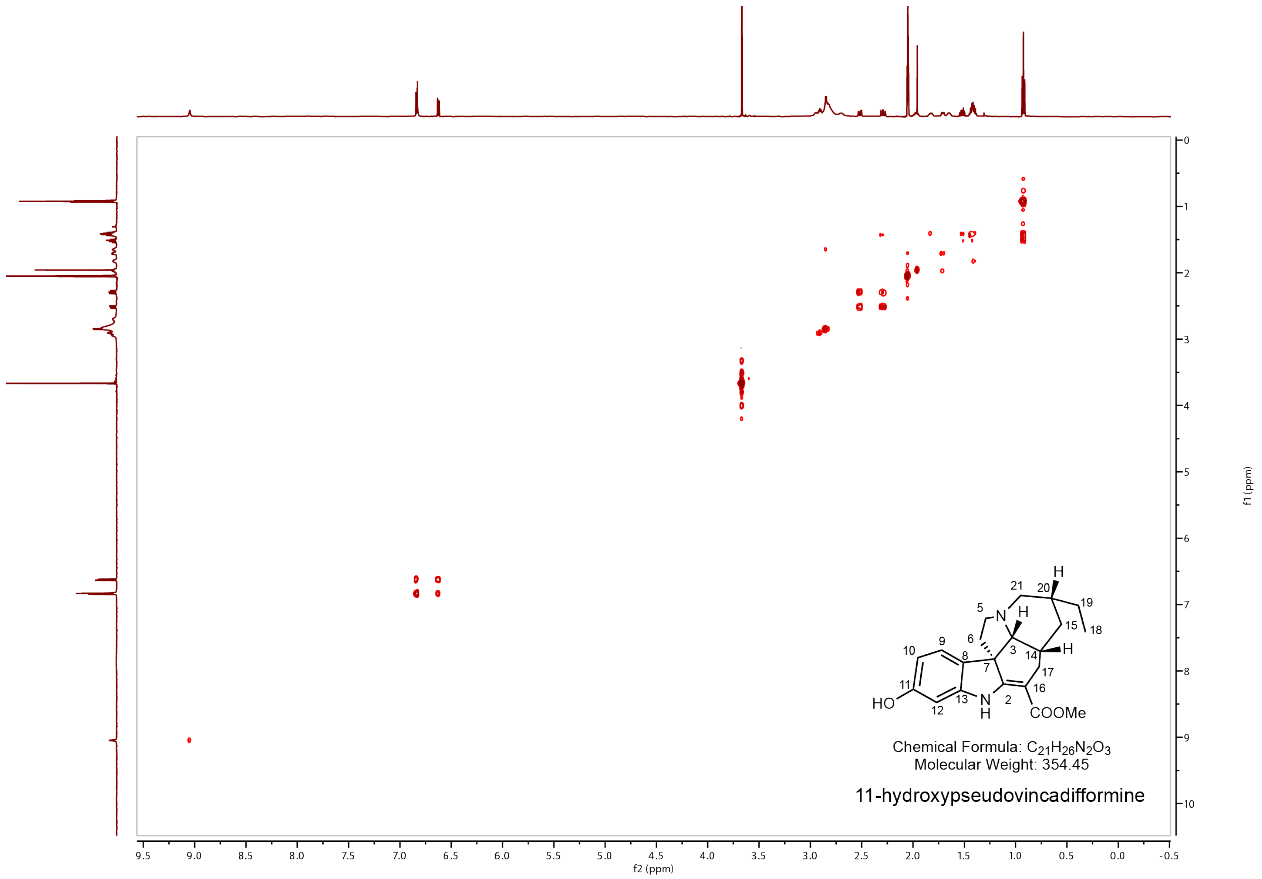


**Supplementary figure 5. COSY NMR spectra for 11-hydroxypseudovincadifformine in acetone-*d6*.**

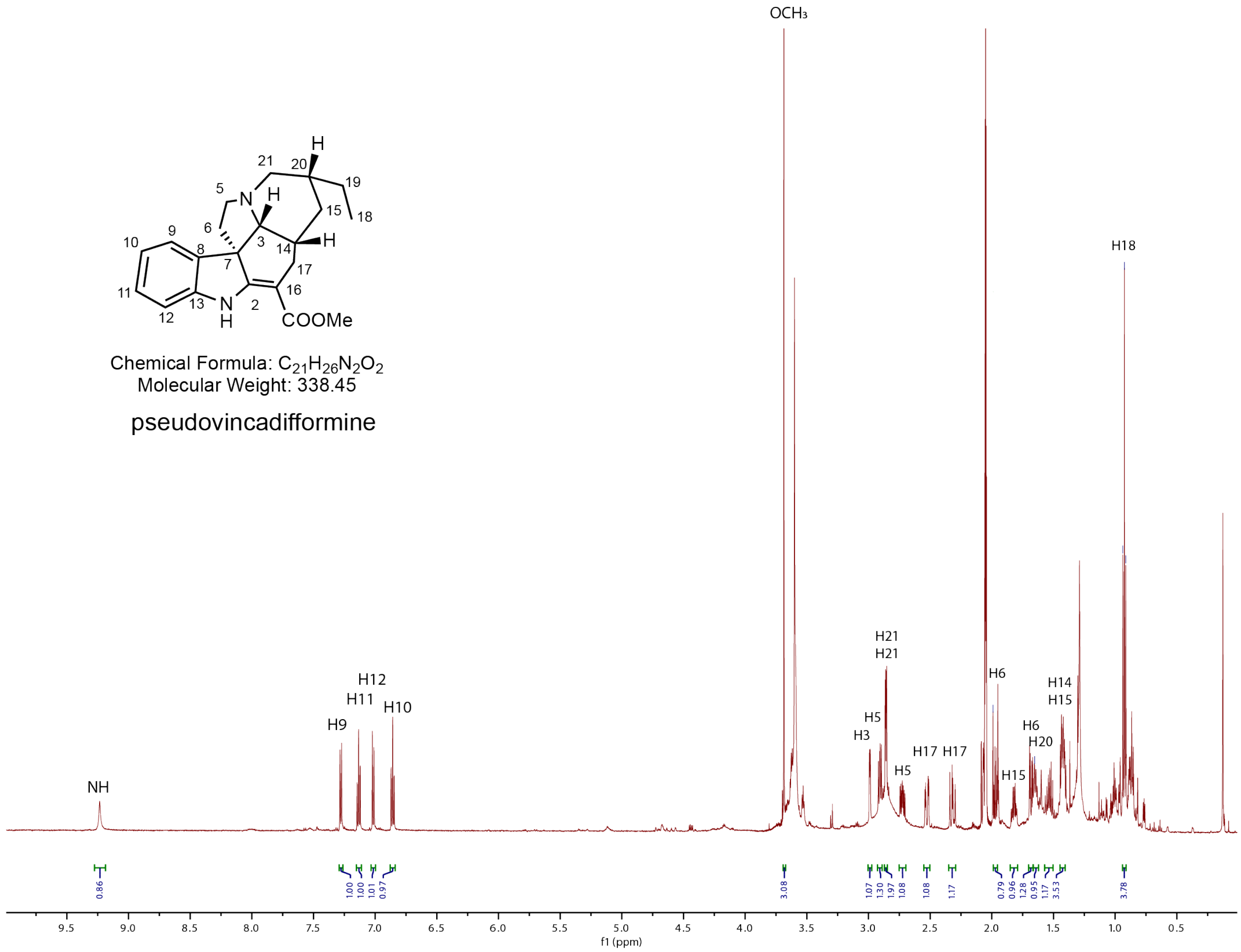


**Supplementary figure 6. $^1$H NMR spectra for pseudovincadifformine in acetone-*d6*.**

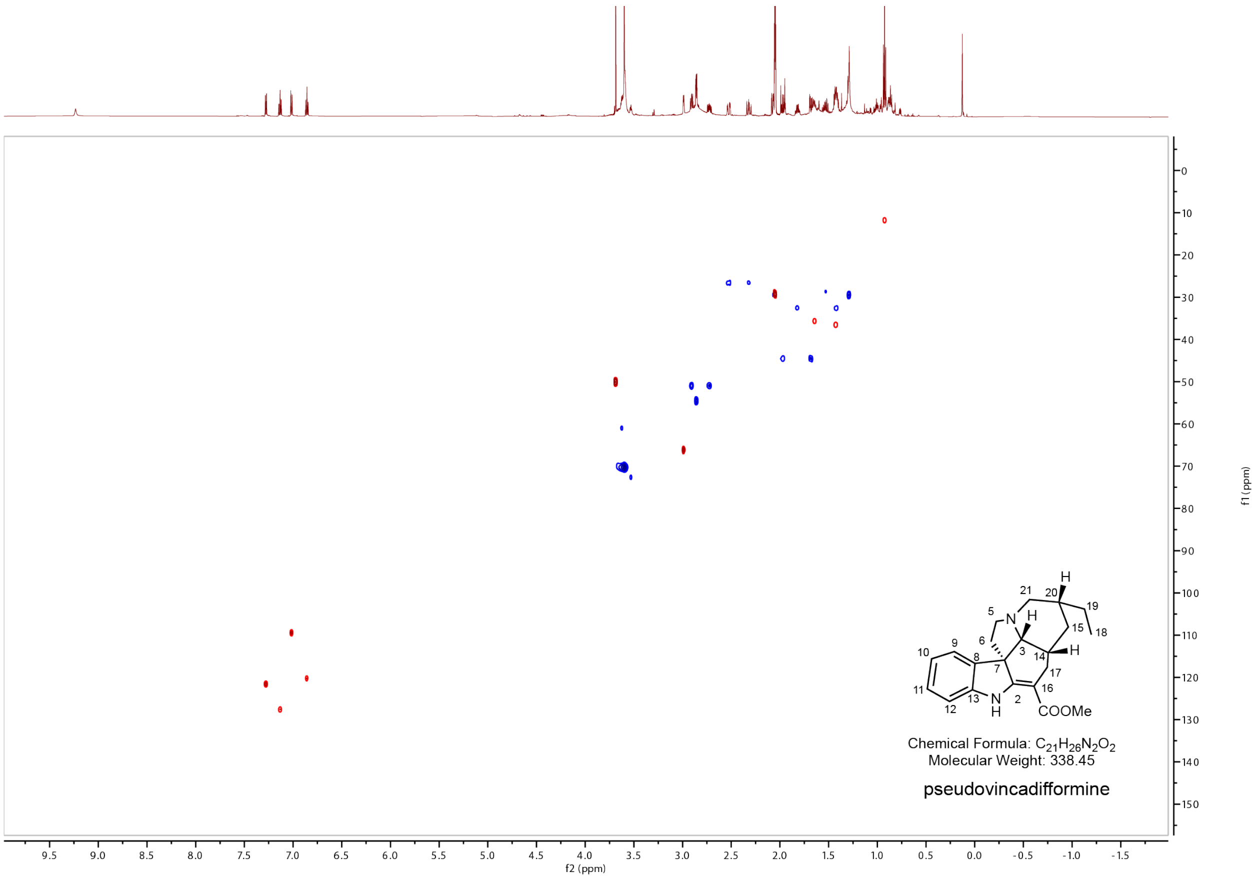


**Supplementary figure 7. HSQC NMR spectra for pseudovincadifformine in acetone-*d6*.**

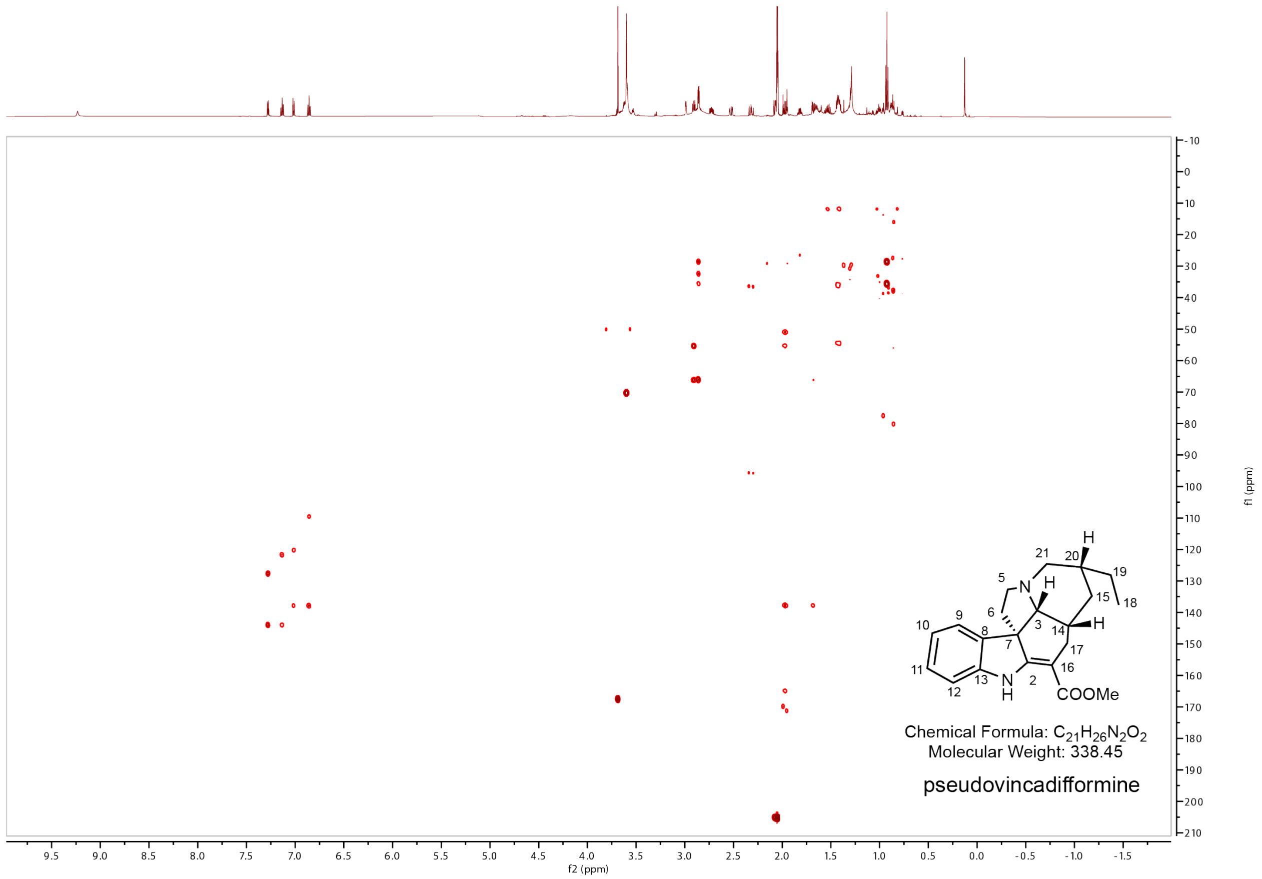


**Supplementary figure 8. HMBC NMR spectra for pseudovincadifformine in acetone-*d6*.**

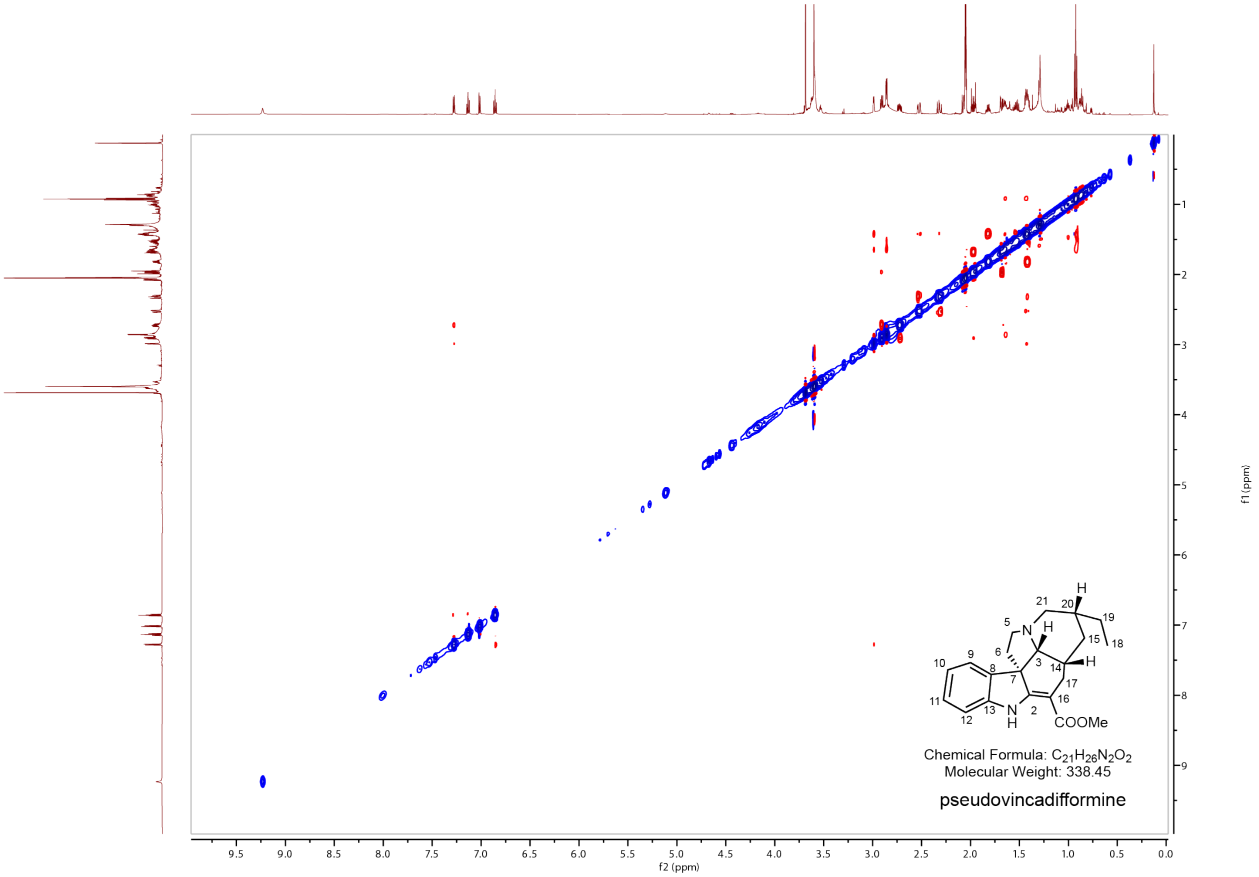


**Supplementary figure 9. NOSEY NMR spectra for pseudovincadifformine in acetone-*d6*.**

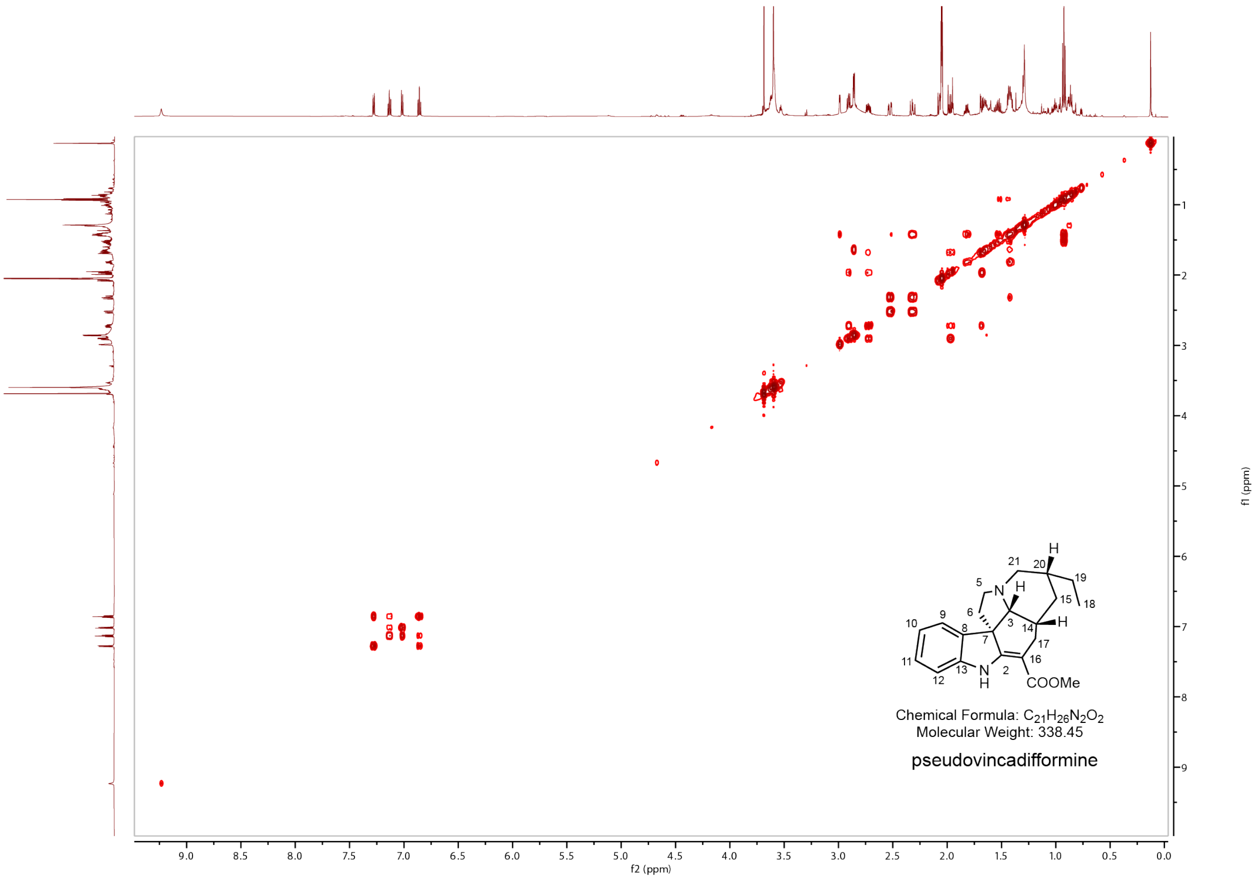


**Supplementary figure 10. COSY NMR spectra for pseudovincadifformine in acetone-*d6*.**

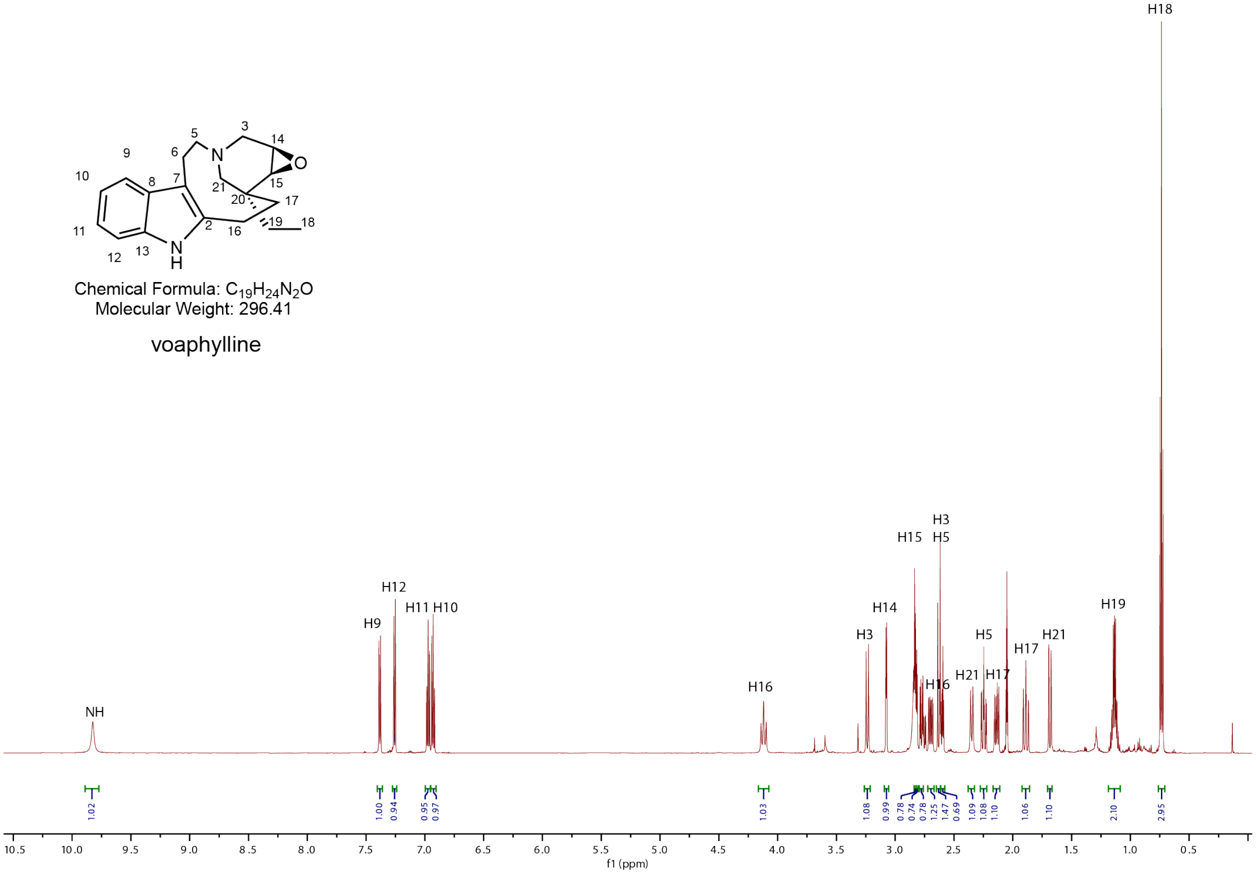


**Supplementary figure 11. $^1$H NMR spectra for voaphylline acetone-*d6*.**

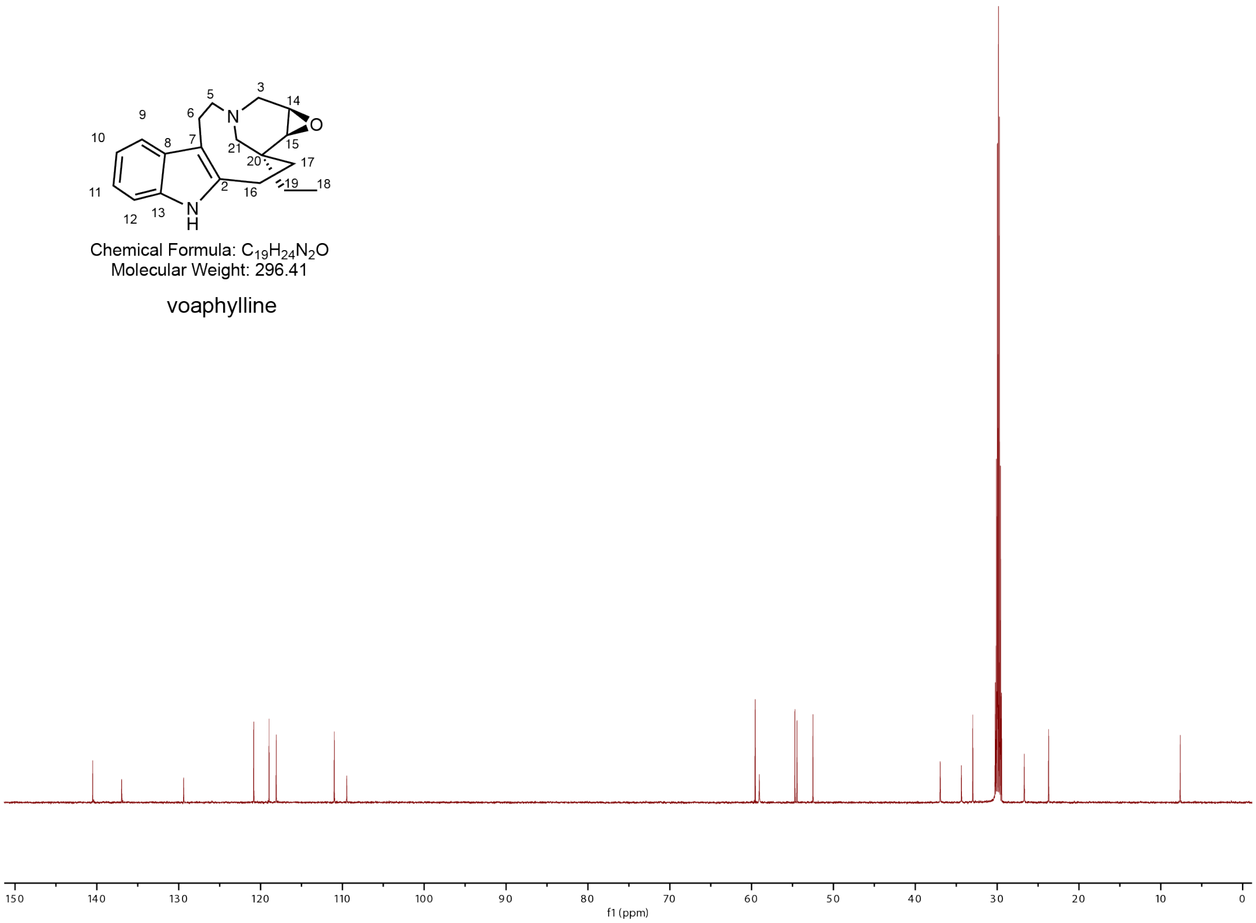


**Supplementary figure 12. $^{13}$C NMR spectra for voaphylline acetone-*d6*.**

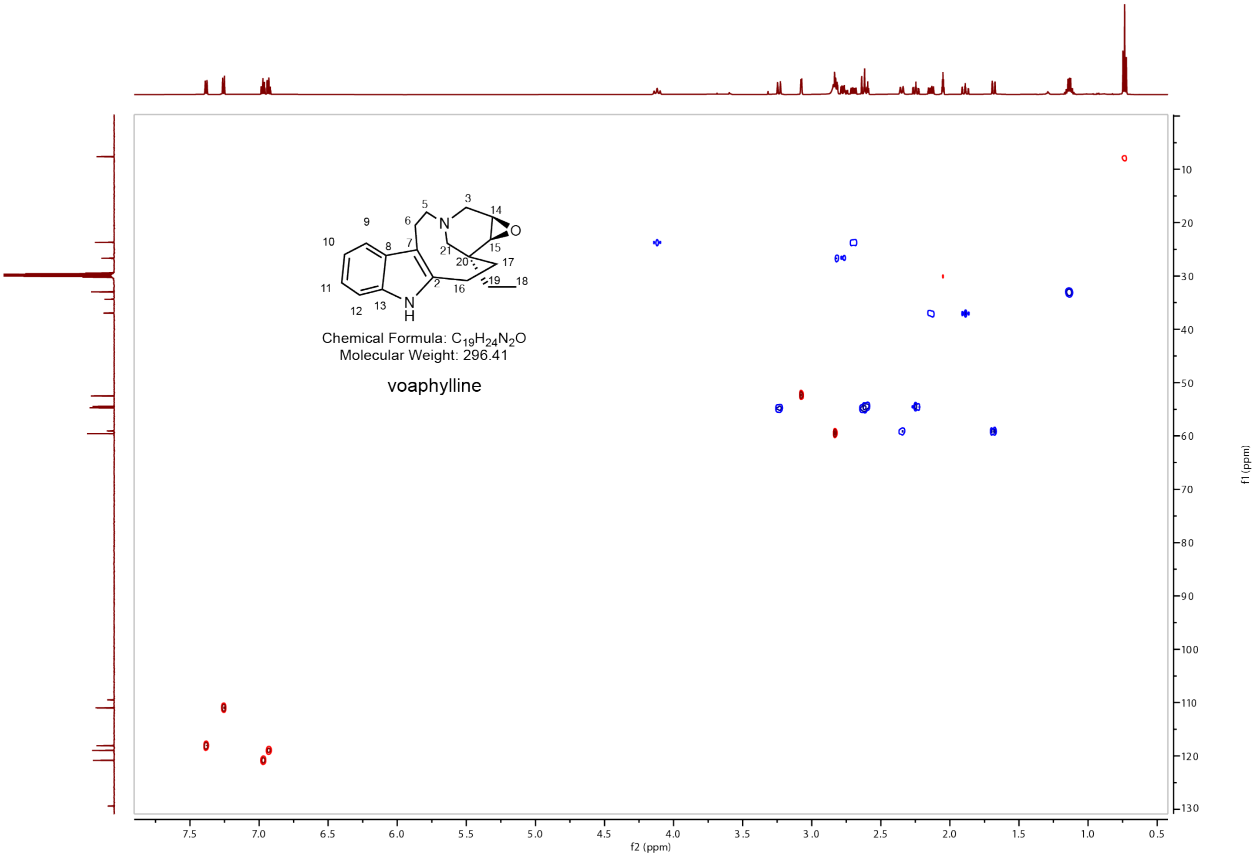


**Supplementary figure 13. HSQC NMR spectra for voaphylline acetone-*d6*.**

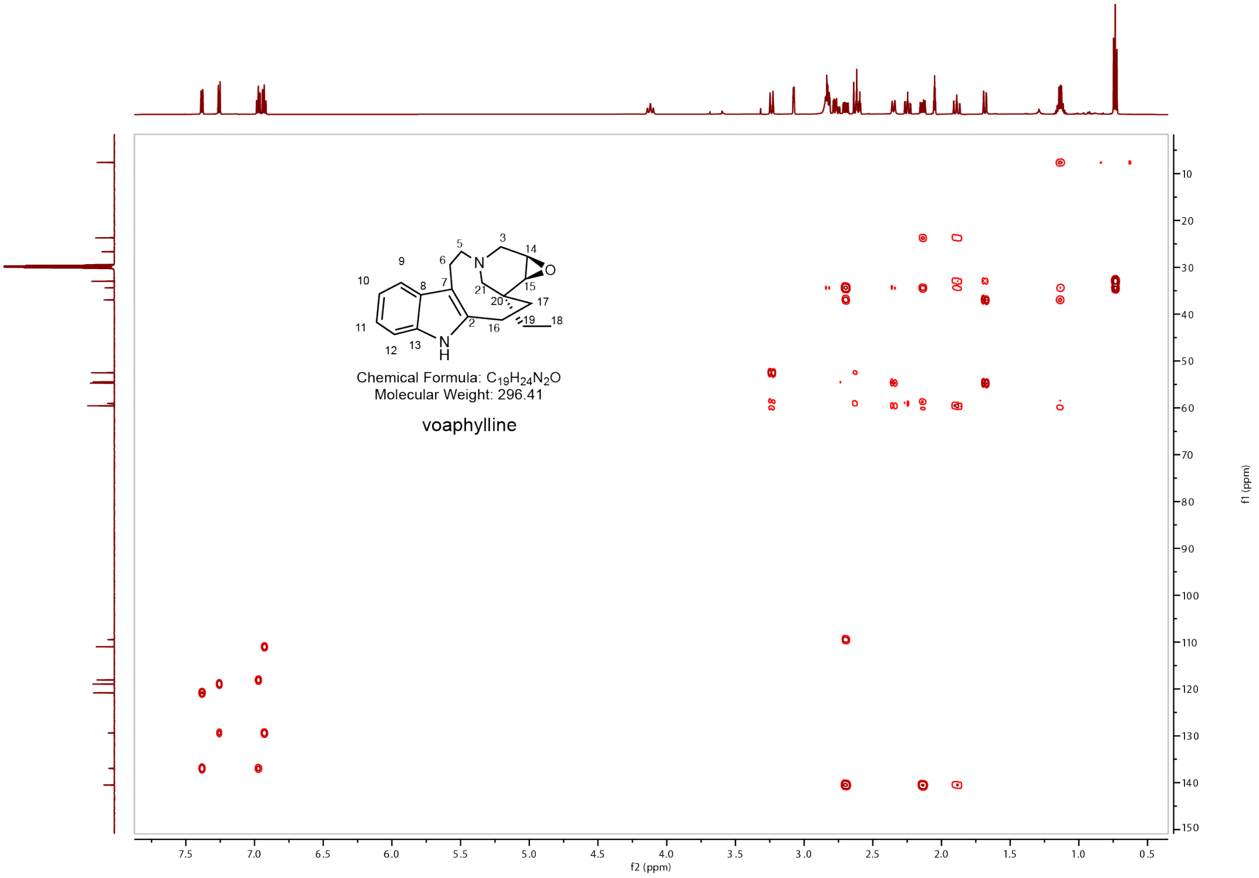


**Supplementary figure 14. HMBC NMR spectra for voaphylline.**

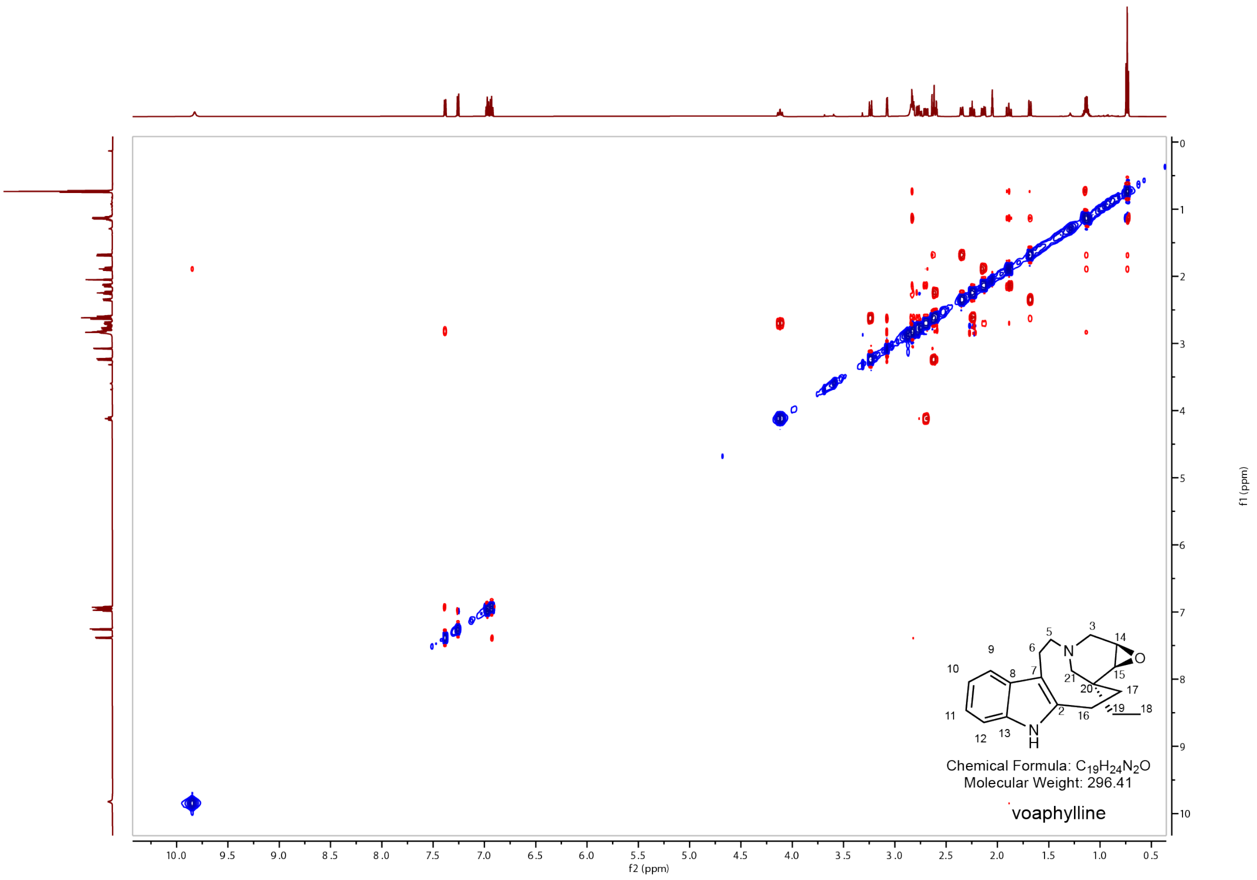


**Supplementary figure 15. NOSEY NMR spectra for voaphylline.**

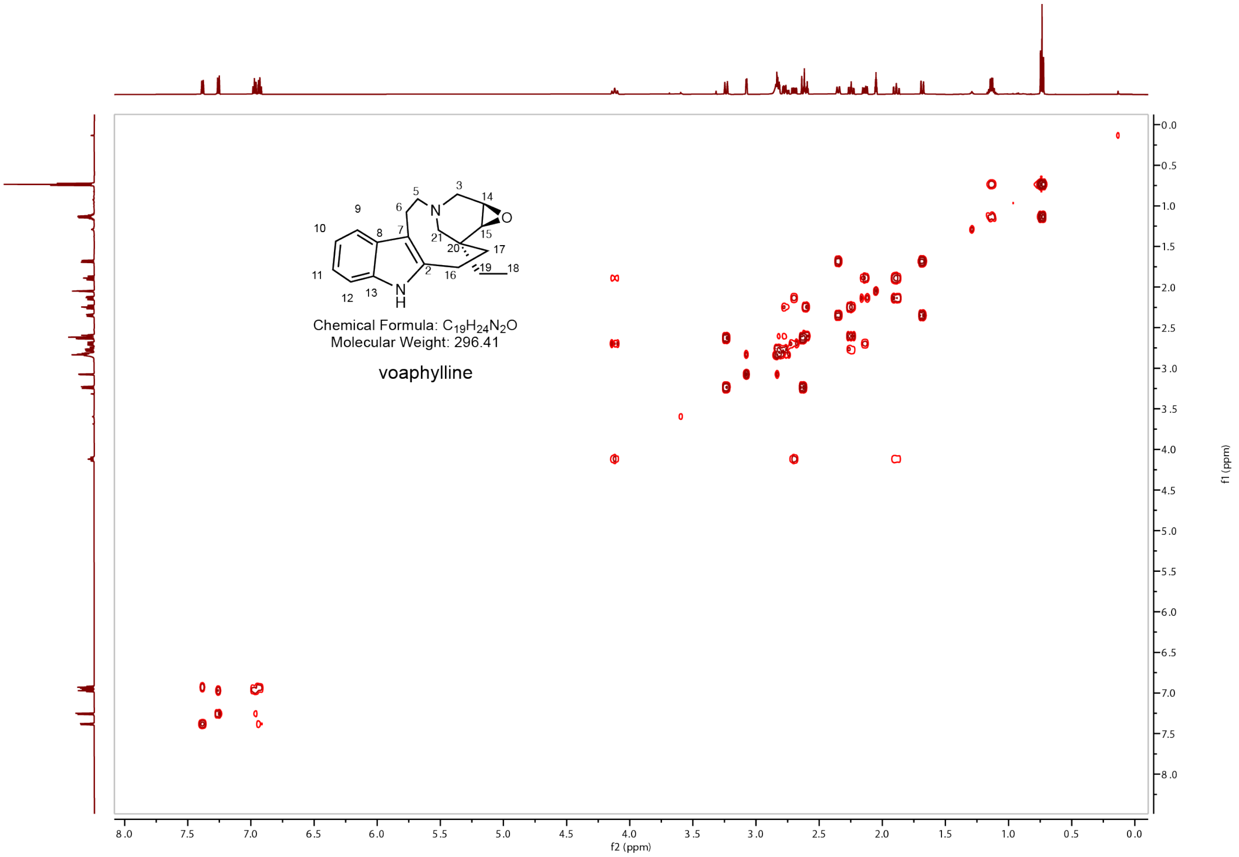


**Supplementary figure 16. COSY NMR spectra for voaphylline in acetone-*d6*.**

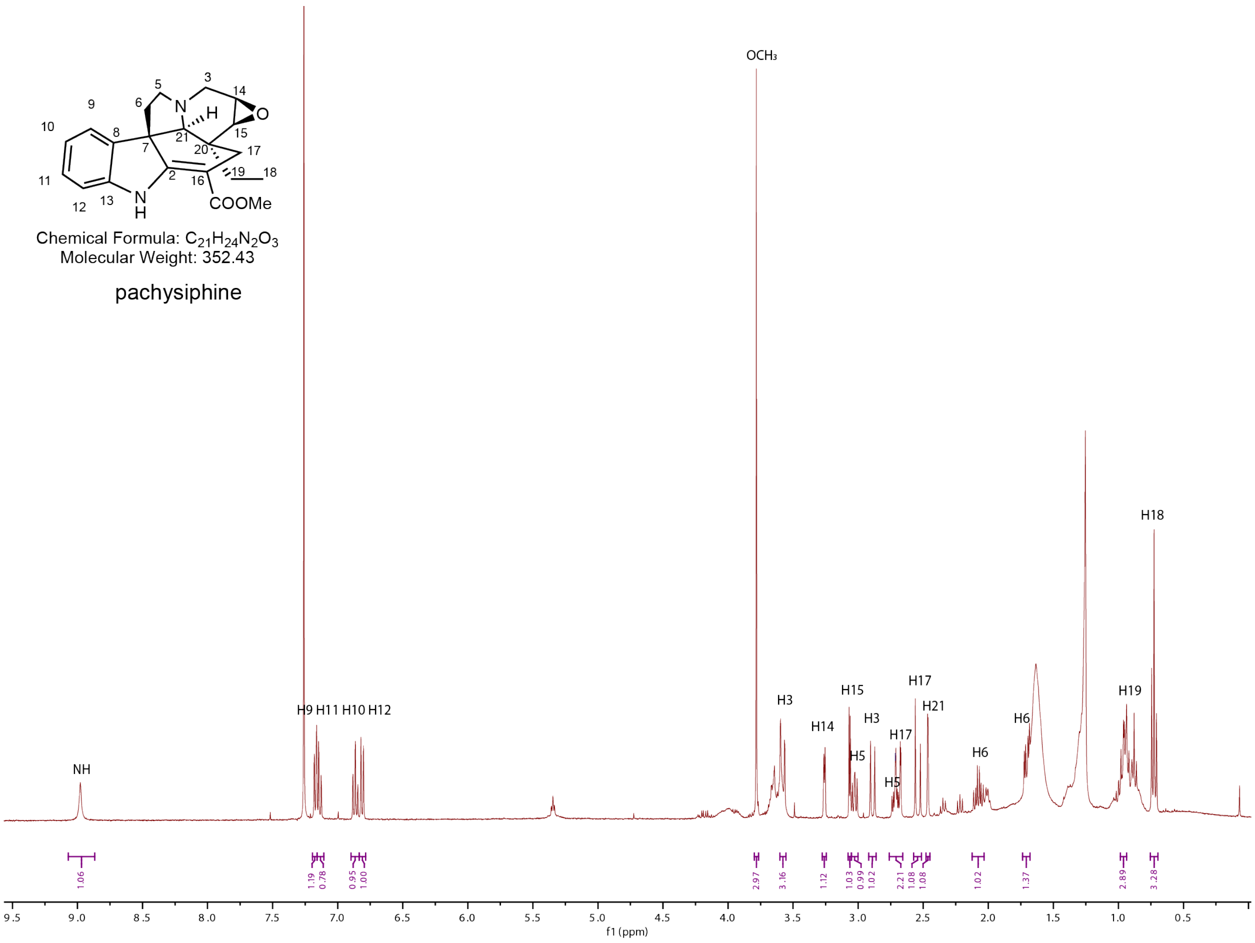


**Supplementary figure 17. $^1$H NMR spectra for pachisiphine in $CDCl_3$.**

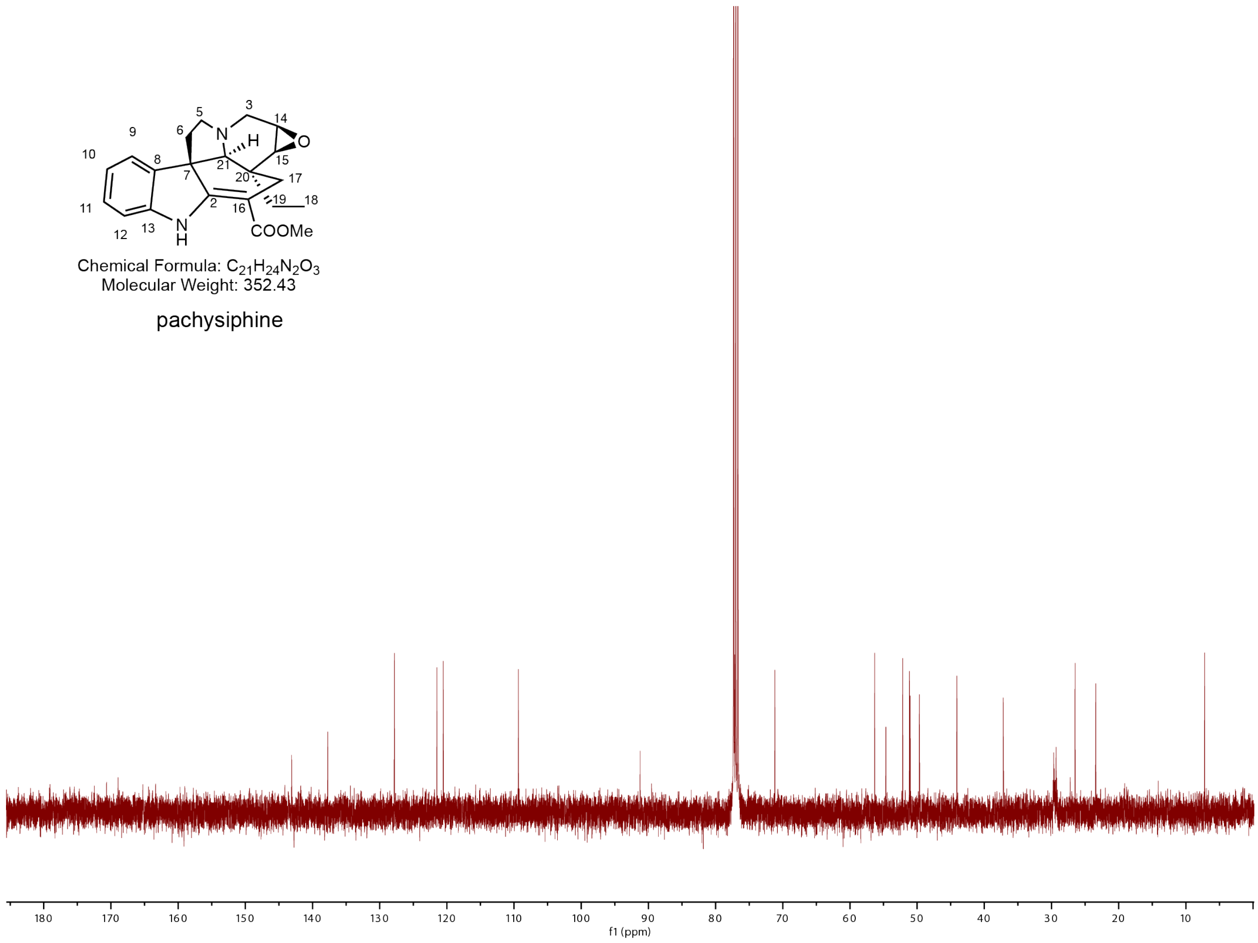


**Supplementary figure 18. $^{13}$C NMR spectra for pachisiphine in $CDCl_3$.**

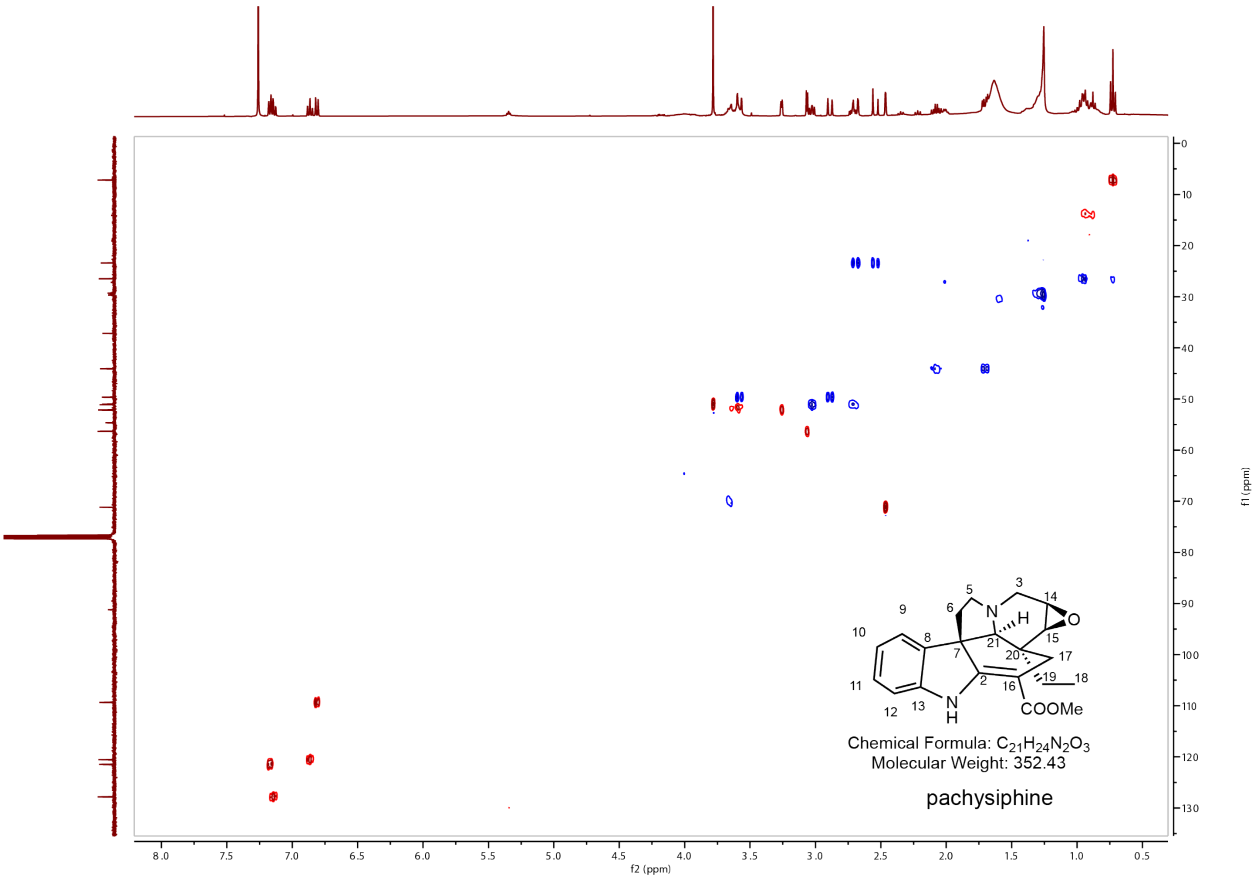


**Supplementary figure 19. HSQC NMR spectra for pachisiphine in $CDCl_3$.**

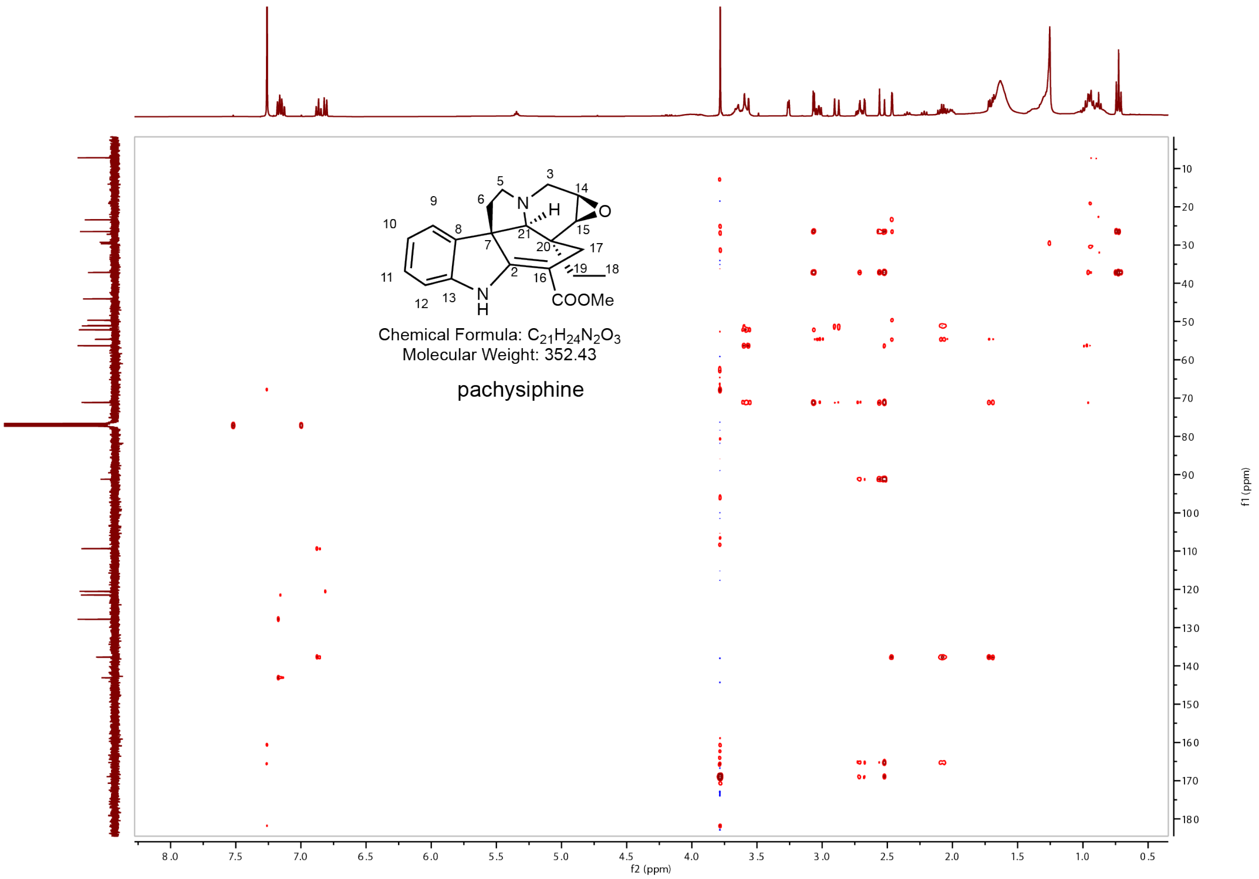


**Supplementary figure 20. HMBC NMR spectra for pachisiphine in $CDCl_3$.**

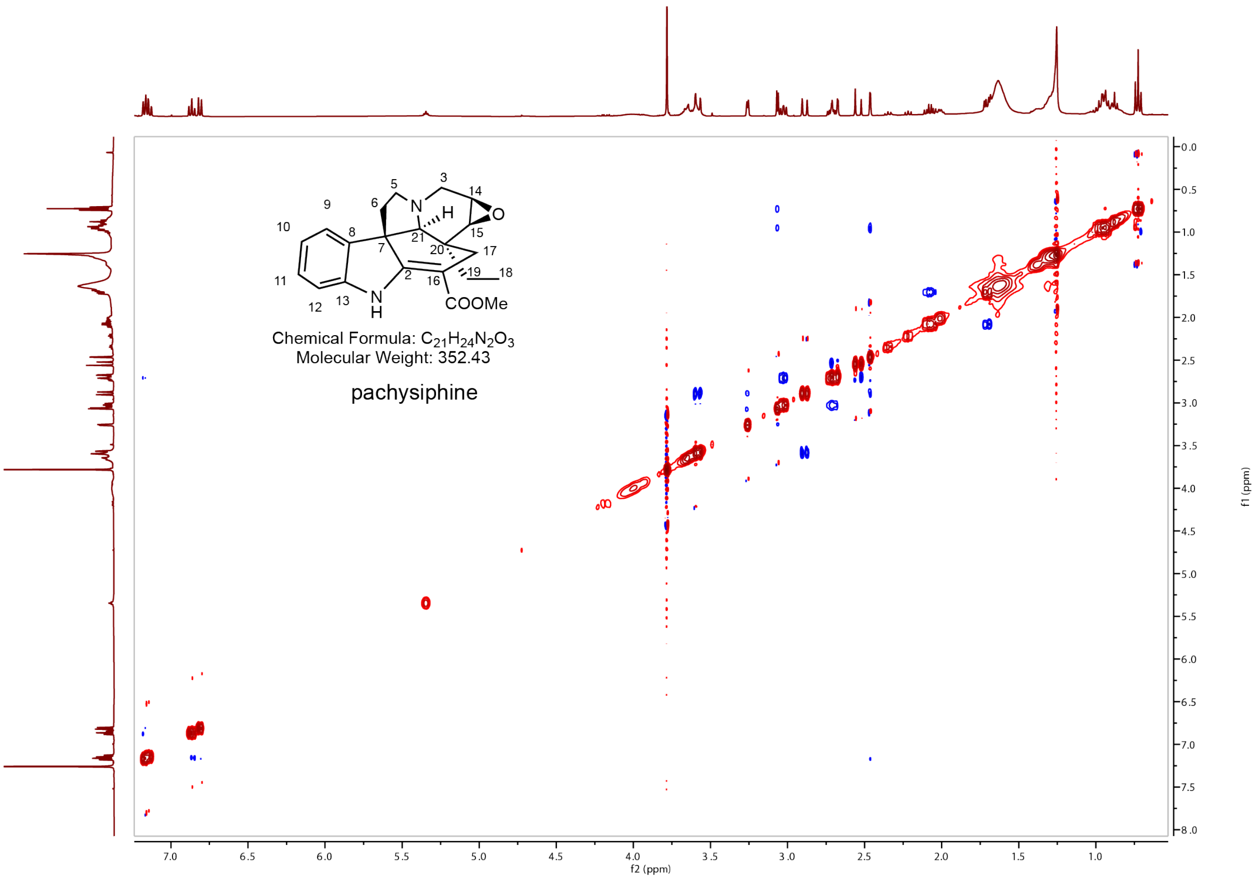


**Supplementary figure 21. NOSEY NMR spectra for pachisiphine in CDCl₃.**

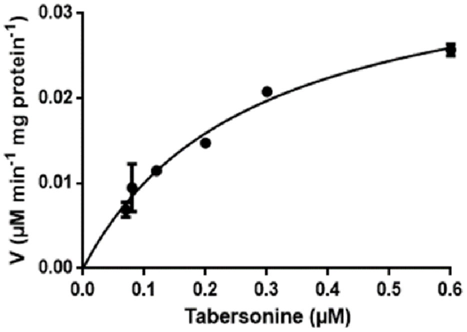


$K_M$ 0.277 ± 0.036 µM

$V_{max}$ 0.0379 ± 0.0024 µM/min/mg protein

**Supplementary figure 22. Saturation kinetics for TliTbE.**

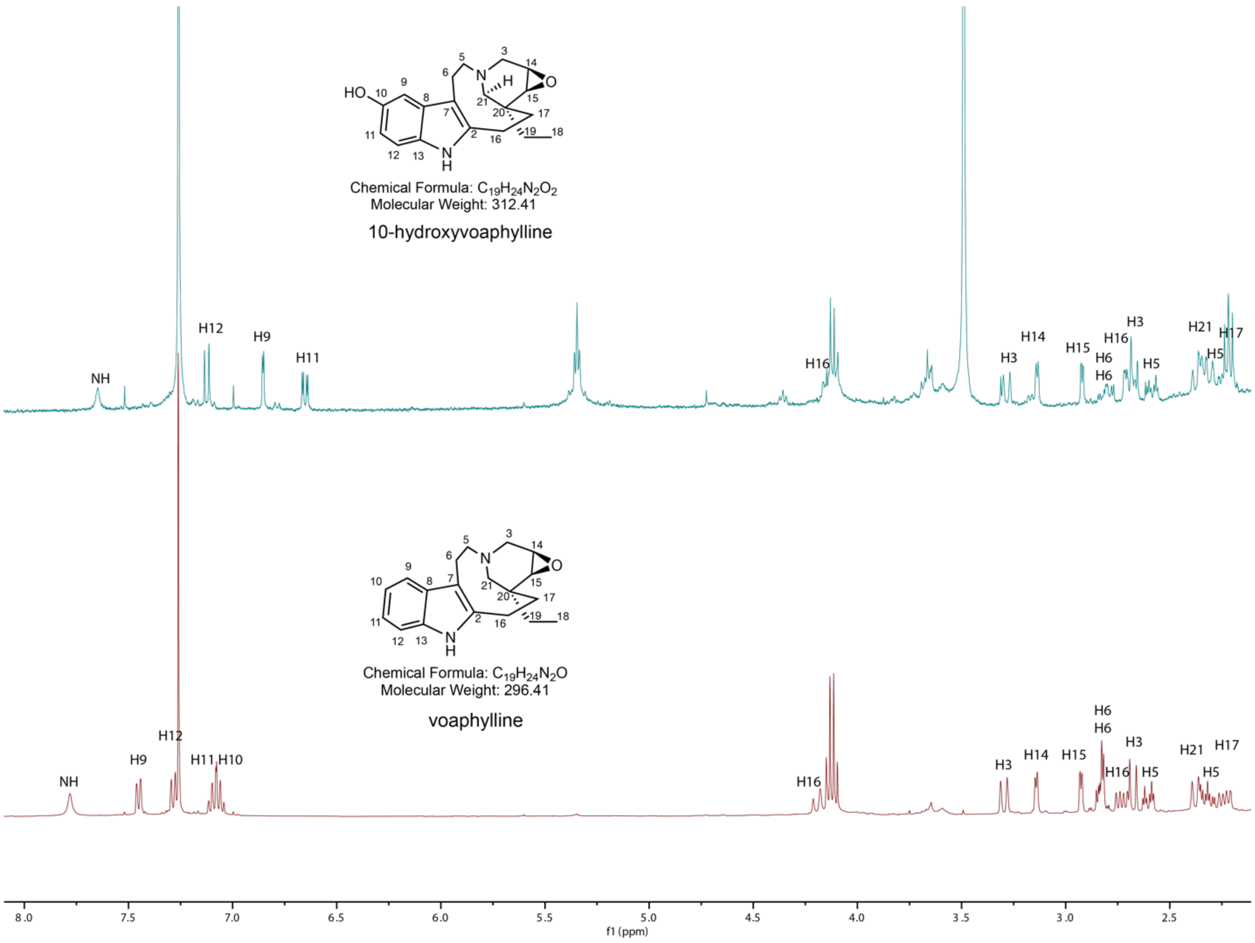


**Supplementary figure 23. 1H NMR spectra for 10-hydroxyvoaphylline and voaphylline in $CDCl_3$.**

**Supplementary data 1. Protein alignments used for phylogenetic studies.**

```
>TliCYP4
----------------------MDILSSILALIFLLAFIANHSLVKKNRSKIKENEKFRVPEPPGAWPFLGHLHLL
AGQ-VPIARVLGAMADK-HGPIFSLRLGSRRALAVSSWEMVKECFTTNDRIFASRPV-MAVAKYIGCDNAIFSLAP
YGPYWRDIRKMVTLELLTNQRLEKLKHVRASELDNLIKNFYS---LCSNDPSPARINLNNWLEYLTFNMILRTLAG
KRFSSCDDGKDDEDWK-IKNAIKKALYLSGVFVVSDVIPS--LEWLDIGGHIKAMKQAGKELDEVLQPWLEEHIQR
RKQSDVEGEA-------DFMDVMLSAYPEN-TIISGHKREDIIKSTTTNLILTASESTAETLVWALSLLLNNRAIL
KLAQDELDVQVGKTRWVEESDISNLKYLQAIVKETLRLYPPGPLAGPREATEDCNIGGYFVPKGTRLIVNLWKLHR
DPRVWTD-PLEFKPERFIDSHTNVSYKG-QSYEYIPFSSGRRMCPAVNYGLQVVHTTLARLIHSFDISTPMDAA--
--VDMSEGLGIALPKVKPLEVVLTPRLPLELYQKL---------
>RsVH
----------------------MDLLQILLAIAGLLAILLLQKQWRTKTSPGAKAGRKLPPEPAGAWPVIGHLHKL
GGP-NPIYRNLAEWSDK-YGPVMTLKLGMQNAVVVSDREAIKECFTTNDKALADRPP-SSIGLHLGFNYAAIGAAP
YGPYWRDMRKLVLLEVLSSRRLEMLRNVRISEIGTSIKELYSNIIRSSGGSGPAKVVISHWIEQLTLNYILRTIAG
RRFS---DDSSKDAQY-VKGVINDFMYFAGQFVVSDVIPIPLLRWLDPQGHLKGMKRVAKEVDTMCEAWIQEHVQR
RMRE---------------------------------------------------KPGPGQEQDFIDVLLNNRDVM
RKAQEEIDNHVGKERWVDETDLKHLVYLQAIVKEGLRLYPPGPLGAPHRAIEDCQVGGYFIPKGTQLLVNVWKLHR
DPRVWSE-PEKFMPERFLTRQAEVDVFG-HHFELLPFGSGRRACPGITFAVQVMHLTVARLLQGFDMTTPSNLP--
--VDMTEGPGVTMPKAHPVEVLMMPRLPSALYEP----------
>Snv10H
------------------MEFSLLYIHTAILGLISLFLILHFVFWRLKSAKGGSAKNSLPPEAGGAWPIIGHLHLL
SGS-KLLHITLGSLADK-CGPAFIIRLGVRQALVVSDWELAKELFTANDVAISSRPK-LLAFESMSYDFAMFGFSP
YGAYWRELRKLISVELLSTRRLELLKHIRVSETEISVKELYNLWKDKKNGSGHVLVEMKQWFGDLNLNVILRMVAG
KRYFGTIDAKDQEEARRCQKAMSGFFHFTGLFLVADGFPF--LRWMDLGGYEKKIKASAKEMDLIAEEWLQEHRRK
RESGDVASEQ-------DFMDLMLSLLEDV-DLPG-YDPDTITKATCINLILGGADTNTVMLTWTLSLLMNHPHIL
RKAQEELDIQVGKERRVNESDIANLEYLHAIVKETLRLYPASRLGGPREFSEDCTLGGYHVTKGTSLILNLWKLQR
DPRIWSN-PSEFRPERFLTTHKDLDVKG-RYFELIPFGAGRRSCPGTAFGLQMLPFVLANLLHAFDIST--DEK--
--TDMTESPGLTTSKATPLDVLISPRLSPNLY------------
>CrT16H2
------------------------MELYYFSTFAFLLFCFILAKTLKKSGQ----SNLKLPLGPPPIPILGNAHQL
IGG--HTHHILRDLAKK-YGPLMHLKTGEVSTIVASSPEIAEEMFKTHDVLFADRPSNIVAFKILSYDYSDVVISP
YGNYWRQLRKISMMELFSQRSVQSFRSIREEEVLNFIKSIG--------SREGTKINLSKEISLLIYGITTRAAFG
EK--------NKNTEEFIRLLDQLTVAV-AEPNIADMFPS-INFLKLISRSKYKIEKIHKNFDAIVQTILNHHKDR
LANHKSSSHEENGEQNKDLVDVLLNIQQRG--DFDTPLGDRSVKAVIFNIFSAGTETSSTTVDWAMCEMIKNPTIM
KKAQEEVRKVYNEEGNVNETKLHQLKYLKAVIKETLRLHPPVPLLLPRECREQCEIKGYTIPSKSRVIVNAWAIGR
DPNYWIE-PENFNPERFLESEVDFKG---NSFEYLPFGGGRRICPGITFALANIELPLAQLLFHFDWKLASDETNI
DKLDMTESRGVTVRREDDLCLIPFPYSASSLKGKY---------
>CrTEX1
-------------------------MEFVVSLFAFVVSCFILLKVAKNSKNPKRNTNLELPPGPKQLPIIGNLHQL
GGG--LAHHVLRNLGKQ-YGPLMHLKIGELSTIVVSSTEIAKEVFKTHDIHFSNRPSHILVFKIVSYDYKDIVLSQ
YGKYWRELRKVCNLELLSPNRVQSFRSIREDAVLNMMKSIS--------SNDGKVVNLSEMILSLIYGITARAAFG
VW--------SKKHEEFIRLESEIQRLA-TTFVLADMFPS-IKFLGALSGLRYKVEKVHKKVDDILEGILKEHRRQ
NNNMTEENGK------KDLVDVLLNIQKNG--DMETPFTDQHIKAIIFDMFSAGTLTSTIAVDWAMAEMMKNPSVL
KRAQDEVRNVYNGIGNVDESKLDELKYLQAVIKETLRIHPGTPIVH-RETREECEINGYRIPAKARVMVNAWAISR
DPNYWPE-PDIFKPERFLGSEVDFKG---THFEYIPFGAGRRICPGISYAIANVQLPLAQLLYHFEWKLPGGMK-P
EELDMTEILGTAAQRKENLLLIPNSHSCSSLKQV----------
>TliC10H
-------------------------MELIVSLFAFLIFCLISVKLSKKSNA----AKLKLPPGPRQLPIIGNMHQM
LGG--QPHHVLRDLAKK-YGPLMHLKFGEVSTIVVSSPEMVEEVFKKHDVLFADRPSNLIAFRISSYDFLDIFASP
YGDYWRQLRKICMMELLSAKRVQSFRSIREEEVLKFIRSIS--------VQEGSVINLSQKLFSLVFGITARAAFG
RK--------NRYDEQFHHIDDKMVKLS-SGFCIADMYPS-VRLFQVMSGVKRKLEKLQKQLDEILENILNEHKEE
SMKQKFGSGG----AKEDLIDVLLNVHKRG--DFGAPLSNSSIKAVISNMFSAGSDTSSAAIEWAISEMIKNPTIM
KKVQDEVRNVFEETGNVDETRLGELKYLQAVIKETLRLHPSAPLLAPRECREQCEIKGYTIPAKARVFVNAWAIGR
DPMYWTE-SEKFIPERFLGSEIDYKG---NHFEYIPFGGGRRVCPGMSFGLANIQLPLAQLVYHFDWKLSGDLK-P
EELDMTEADKALSERKNKLHVIPMVYSRSFMKK-----------
>TiI10H
```

------------------------MEMLIVSFFAFLIFCLISVKLSTKSKA----DKLKLPPGPRQLPIIGNMHQM
LGG--LPHCVLRDLAKK-YGPLMHLKFGEVSTIVVSSPEMAKEVFQKHDILFADRPSNLIAFKISSYDFLDIFASP
YSDYWRQLRKICTMELLSSKRVQSFRSIREGEVLNLIRWIS--------VQEGSVINLSKKLFSLIFGIITRAAFG
KK--------NRYEEQFHHLDEQMTKLA-SGFCIADMYPS-VRLFQVMSGVKPKLENLQKQLDEILGNILEEHKEE
SMKQKLGGGE----AKEDLIDVLLNVHKRG--DFGAALSNSSIKAVICNMFSAGSDTSSSALEWAISEMIKHPTTM
KKAQDEVRNVFQETGNVDETKLGELKYLQAVIKETLRVHPSAPLLAPRECREQCEINGYTIPAKARVFVNAWAIGR
DPMYWNE-PEKFIPERFLGSEIDYQG---NHYEYVPFGGGRRVCPGMSFGLANIALPLAQLIYHFDWKLPGDLK-P
EELDMSEDVSALSARKNNLYIIPMVYSRSFMKKEITAEVMMDEA
>TliCYP10
-------------------------MEVVFLICAFLFFCFMLVKLAKKS------TGLKLPPGPRPLPVIGSMHQL
YGG--LIHHKLRDLAKN-YGPVMHLQLGEILTIVITSPEAAREVYKTHDIIFSQRPSCFESFRAISYDFTDIIFSP
YGNYWRELRKICTMELLSPKRVRTFRWIREEEVLNLIRSIS--------SQEGSVVNLSHKIHSLSYCITSRAAFG
NR--------NKDQEKFQRLVQEIIKVA-SGFNLSDMFPS-AKWLRVLSGMEHKLQMVQKQVDEIVQKILNEHKVE
MMKRETQSRE----AKEDLVDVLLNIQRSG--HFEVPLTDDNLKAVIFNIFGAGSDTSSTEVEWAISETIKNPRIL
KRAQDEVRNVFNERGNVDESRIHQLKYLQAIVKETLRLHPSAPLLLPRECSVRCNIHGYDIPAKARIFVNAWAVAR
DPKFWNE-AEKFNPERFLDSEVDYRG---NDFEYVPFGAGRRICPGISYALPNILLPLAQLLFHFDWKHPGVLQ-P
EDLDMDELMGVTVRRKNDLQLIPMLYHSSSLKK-----------
>GsR11H1
-------------------------MELIIFIAVFLLFCTLVVKIVKKSKP------VKLPPGPKPLPIIGNIHQL
SGG--LPHHILADLAKK-YGSLMHLKLGEVSHIVVSSAEVAKEVMNQHDILFSSRPE-LLAVRIIDYNCTDITFSP
YGDYWWQLRKICTVELLSAKRVQTFRSLREEEVFNMIKLVF--------HKKGSVMNLSKMVFSVTYSIIAQAAFG
KR--------SKYQEEFLVLMEEMMNVS-GGFSIADLYPS-IKMLEVITGMRQRLEKIHKKKDIILGNILEEHKVK
RIKSD--ARGDQ---REDLIDVLINVQSSV--EFGAPLTDDNIKAVISEIYSAGGESSATTVIWTISEMIRNPGVL
KRAQDEVRQVFDGKGNVDESGLHELQYLQAAIKETMRLHPALPLLLQRECRESCEIDGYEIPVKTRVIINAWAIGR
DPMHWNE-PEKFNPDRFLGSQIDFRG---KDFSYIPFGAGRRMCPGITFALPNVQLPLAQLLFHFDWELPGGLE-L
EKLDMTENFGLAVGRKHDLDLVPTPYSGSSLK------------
>TliCYP7
-------------------------MEIIFLLCASLLFIFVFVKTAKKSKP------MKLPPGPRTLPIIGNLHQL
SGP--LLHHILADLAKK-HGPLMHLKLGEISTMVISSPEVAKEVMNDQDVIFAHRPY-TLAAKILHYNRSDIAFSP
YGDYWRQLRKICMVELLSAKRVRGFRSIREAEVQNMIRSVS--------LHKGSAANLSKMFFSLTYSIIAQAAFG
KK--------TKYQEEFVRVMEDVMKLI-GGFSIADMYPS-FTIVEVISGMRQKLQKTHQKIDRIMGTILDEHKEE
SRK----TRGKE---EKDLVDVLLDIQSSG--DFEVPLTDDSIKAVIFDIFGAGGETSATTSIWAMSEMIKNPRVL
RRAQEEVRKVFSDRGNVDESSLHELKYLQAVIKETLRLHPPAPLLLPRECSEQCKVYGYDIPAKAILFINLRAIGR
DPMHWTE-PETFNPERFLDSEFDFRG---TNFSYIPFGAGRRICPGISFGLPNVELPLAQLLYHFDWKLPGELQ-P
EELDMVEKFALTVGRENDLELIPIPYSRSYLQ------------
>TliPs18H
---------------------MEFFTALCGVFLVLYVFLIFINSYKSSKSS------LKLPPGPKPWPIIGNMHQL
RGS--LTHRVLGDLAKT-HGSLMHLQLGGVSAVVVSSPEAAQQFMKTHDTTFAYRPQ-LLASRIMTYDGTDIVFAP
LGDYWRQLRKICMMELLSPKRVQSFRPIREEEMLNLIRSIS--------LNKGSAINLGRKIFSYSYGVAARAAFG
KR--------NKYQEEFTSTVEEALKLS-TGFNLADFYPS-VKFIQVIGKINPKLKRLHKTLDAILENIVNDHREK
RMKIEPSSDGET---KDDLVDVLLNIQKSG--DFGAPLTDNNIKAVILDIFSGGSETSSTTVEWAMSEMVKNPQVM
KRAQDEVRKVFNGKGNVDESGLHELKYLQAVVKETLRLHPSAPLLLPRECSEPCKIDGYDIAIMTRVIINAWAIGR
DSRYWDE-AEQFIPERFLDCAIDYHG---RDFKYIPFGAGRRICPGISFALPNIELPLAQLLYHFDWKLPDELK-P
EELDMTETFSVTVRRRHHLCLVSIPYSHSSHV------------
>CrT3O
-----------------------MEFHESSPFVFITRGFIFIAISIAVLRRIISKKTKTLPPGPWKLPLIGNLHQF
LG--SVPYQILRDLAQK-NGPLMHLQLGEVSAIVAASPQMAKEITKTLDLQFADRPV-IQALRIVTYDYLDISFNA
YGKYWRQLRKIFVQELLTSKRVRSFCSIREDEFSNLVKTIN--------SANGKSINLSKLMTSCTNSIINKVAFG
KV--------RYEREVFIDLINQILALA-GGFKLVDLFPS-YKILHVLEGTERKLWEIRGKIDKILDKVIDEHREN
SSRTGKGNGCNG---QEDIVDILLRIEEGGDLDLDIPFGNNNIKALLFDIIAGGTETSSTAVDWAMSEMMRNPHVM
SKAQKEIREAFNGKEKIEENDIQNLKYLKLVIQETLRLHPPAPLLMR-QCREKCEIGGYHIPVGTKAFINVWAIGR
DPAYWPN-PESFIPERFDDNTYEFTKSEHHAFEYLPFGAGRRMCPGISFGLANVELPLALLLYHFNWQLPDGST--
-TLDMTEATGLAARRKYDLQLIATSYA-----------------
>CrV19H
-----------------------MELDECSPSIFIIS-FIFIAISIAILRRIRPKKTKALPPGPWKLPLIGNLHQF
ISRDSLPYKILRDLAQK-HGPLMHLQLGEVSAVVASSPEMAKVITRTKDLEFADKPA-IRAIRIVTYDYLDIAFNS
YGKYWREMRKIFVQELLTPKRVRSFWSAREDVFSNLVKTIN--------SANGKSINLTKLISSTTNSIINRVALG

NV--------PYEREIFMELIKQLLTAA-GGFKLVDLFPS-YKIIHVLEGTERKLWKILGKIDKILDKVIDEHREN
LLRTGKGSGENG---QEDIVDILLKIEDGGELDHDIPFGNNNIKALLFDIISGGSDTSSTTIDWAMSEMMKNPQVM
SKAQKEIREAFNGKKKIDENDVQNLKYLKSVIQETLRLHPPAAFLMR-QCREECEIGGYHIPVGTKVFINIWAMGR
DPEHWPN-PESFIPERFENIPYDFTG-SEHQLATFPFGSGRRICPGISFGLANVELSLALLLYHFNWQLPDSST--
-DLDMTEAIGIAARRKYDLHLIPTSYM-----------------
>TliCYP6
----------------------MELLYLFDFITFFLFLAILFATVKKWRRNKKFSPTKKLPPGPPKLPIIGNLHQL
RG--KLPHHAFTDLARKKYGPLMHVQLGEVSAVIVSSPRFAREFMKTHDSVFADRWK-ILVWWIIGYDGLDFAFAP
YTDYWRQMRKICARDFLNNKNVRLFSSIRQDEFSELVAFVH--------SKGGKPIDFTEKILSCTNAIITRAAFG
NA--------FSDRKTFTELLNEIFAAG-GGFDFADLFPS-LRFLTFLLGKKRKLLRMRNRMDSILDTMINEHIEN
---WASAKETNQ---QENIIDVLLRARERG--NTQIPITNNSIKAVLFDMFAGGTETSATVVEWTMSEMLKKPEVM
AKAQTEIRQAMKGKKTVQEDDLPKFKYLNLVVKETLRLHPPLAFLLPRESREDCEIDGYYIPSGTKILVNVWAFGR
DGEYWPD-PESFIPERFENNPIDFKG---NDFEYLPFGAGRRICPGISFGVANVALSLALLLYHFNWNLPDG-D--
-HLDMEEAMGLLATRKNHLRLVATPYDVSM--------------
>TliTbE
-----------------------MELQNLPFNFFAFFVFAFTFLTLVKVWKKSSQEKQNLPPGPWKLPLLGNLHNL
LMG-SLPHHTLRDLARK-HGPLMHLKLGEVNALIVSSPRMAKEVMKTHDLAFANRPV-TLAGKIVCYDYSDIAFSP
YGDYWRQMRKICVLELFSSKCVRSFEPIRKDEGSRLIATLQ--------ASAGKPINLTERISLYTTSMVCRAAFG
KV--------NKGQNKFSQLVKDASEVA-GGFDPADLFPS-YKFLHVLGSTMSKLLKIHGQIDGILEEMVNEHKKN
HAMSKKGNGEYG---EEDIIDILLRIKEGG--DLQFPFTDKNVKGIIFDIFGAGTETSSSVVDWAMAELMRNPKMM
AKAQNEVREAFKGKMTIDETDVRGLSYVKSVIKEALRLHPPVPLLIPRESREPCQIDGYDIPLKTRVFINAWAISR
DKEYWQD-PESFIPERFEGSSVDFTG---TNHEFIPFGAGRRICPGMTFGLANVDFLMALLLYHFDWKLPPGQN--
-DVDMSETIGIAATRKNGLFLVPSPYDIPTLDKSC---------
>RsSBE2
---------------------MEIMNFSLNSPVFLLLSFFFLLMVLMQLTGSRKYKGKRLPPGPKKLPIIGNLHQM
VGS--LPHRVLKNLADQ-YGPVMHLQIGELSAIVISSADKAKEVLNTHGILVADRPQ-TTVAKIMLYNSLGATFAP
YGDYLKQLRQIYALELLSPKTVRSFWTIMEDELSTMVTSVK--------AEAGQPIVLHDRMLTYLYATLCRATVG
SV--------CNGRETLIMAAKETSALS-AAIRIEDLFPS-VKILPVVSGLRTRLTNLLKQLDTVLEDIIGEREKK
MFSSNNIQPSTE---EEDMLGVLLLYKNGKGKDTKFRITNNDIKAIIFELILAGTLSSSAIVEWCMSEMIKNPRVM
KKAQDEVRQVLKDKKKVSGSDLAKLEYVKMVIKESVRLHPPAPLLFPREVREDFEMDGMIIPKKSWVIINYWAVGI
DPKIWDD-AERFEPERFSNSPIDFYG---SHFELIPFGAGRRICPGILFGTTNVELLLASFLYHFDWKLPGGMK-P
EELDMNELFGAGCIRENPLCLIPSISVAGN--------------
>SnvGO
--------------------MMQMMEFSFSSPAFFLLLPFLFLLIKPLIS---RKRGPKLPPGPKKLPIVGNLFHM
EGA--LPHLALKKMTDK-YGPICHLKLGELEAVVVSSAELAKEVLNTHAVTFADRPE-TNVSKIVMYNNSGMTFAR
YGDYFKLLRQIYASELLSPRCVRSSTNHMEDELSKFVVKIQ--------AEAGKPIFLLERVKSYLFAVLFDSMIG
GA--------CKCPERYIEAAKELSANS-AAMRLEDFFPS-VTLLPKLSGFNTVLAKLKKKIDDLLDDLISEREKI
QAN----ATGPM---EEHMLDVLLKLRNGSGSETKVPITNEDVKAVVFELMLS-NLSTAATEEWAMSEMMRSPKVF
KKAQDEVRRVFKGKNRICASELHNLEYLKLVIKEALRMHPPAPLLFPRKAREDCEIGGYTIPVGTMVWVNYWAVGR
DPQLWHD-ADKFEPERFSNVPMDFNG---SHSELIPFGAGRRICPGIAYGVTNLELLLSALLYHFDWELPNGKQ-P
EEIDMDEFYGSGCIRKNPLALIPKVVIPCQA-------------
>CrGO
------------------------MEFSFSSPALYIVYFLLFFVVRQLLK---PKSKKKLPPGPRTLPLIGNLHQL
SGP--LPHRTLKNLSDK-HGPLMHVKMGERSAIIVSDARMAKIVLHNNGLAVADRSV-NTVASIMTYNSLGVTFAQ
YGDYLTKLRQIYTLELLSQKKVRSFYSCFEDELDTFVKSIK--------SNVGQPMVLYEKASAYLYATICRTIFG
SV--------CKEKEKMIKIVKKTSLLSGTPLRLEDLFPS-MSIFCRFSKTLNQLRGLLQEMDDILEEIIVEREKA
SEVS---KEAKD---DEDMLSVLLRHKWYNPSGAKFRITNADIKAIIFELILAATLSVADVTEWAMVEILRDPKSL
KKVYEEVRGICKEKKRVTGYDVEKMEFMRLCVKESTRIHPAAPLLVPRECREDFEVDGYTVPKGAWVITNCWAVQM
DPTVWPE-PEKFDPERYIRNPMDFYG---SNFELIPFGTGRRGCPGILYGVTNAEFMLAAMFYHFDWEIADGKK-P
EEIDLTEDFGAGCIMKYPLKLVPHLVND----------------
>TliCYP5
--------------------------MGFQWEIVLSLLLSFFSCVCIIFCLKRKRIHNYRPPGPPGLPVVGNLLQF
DN--TNRHEFLWQLSKK-YGPLMSLKLGRRRALVISSAKMAREALKTHDLVFSSRPS-VTSQQKLSYNGRDVAFSA
YNSYWREMRKICVLHLFSQKRVQSFQPLLEDEVSHFIQKI-T-----KLASSSQQVNINAIAMTLSSTVICRVAFG
KR-----YDENGHEKKRFKEIFLESQAMLAGFFFSDYFPS-LSWVDRLTGMVARLERTFKELDSFYQELIEEHLDP
NRPQMMKE---------DIIDLLIQLKDDQ--NSQFDITWDHIKAMLMDLFIAAADTIGAALIWSMTALMKNSCIL
NKVQAEIRETVGKKGIVDEEDLRKLPYLKAVVKETLRLYPPAPVPAPRETIDSCFIEGYKIEPKTTVYFNLWAIAR

```
DPEYWKN-PNEFIPERFLDSNLDIRG---HDFQVLPFGAGRRGCPGISLGLATVELALANLLYCFDWELPSGTK-A
EDLDTDVLPGLTMHKKNILWLVAKQYEVFMTED-----------
>Snv11H
----------------------MHSAMSFLLLFSLCFLIHCFVFLLIKKKKAKTMDAKTVPPGPKKLPIIGNLHQL
G---KLPHRSLRCLSNE-YGPLMLMQLGSVPALVVSSADTAREVFKRHDLAFSGRPA-FYVAKKLTYDYSDITLAP
YGEYWREVKKILVLELLSAKKVQSFEAIRDEEVARMVN---------FIARSSNPVNLSRLALSLSNNVICRIAFG
KISG---YEDGSEAKSKFEDIFHETEDFFGTVNIADFFPA-LSWINKFNGVETRLKENFKKLDSFLNQVIEEHLDS
RRSKTEKE---------DIVDALLRIQGNP--NETITLSSENIKGILVTVLLGGSDTSTAVLVWTMAELMRNPMVR
RKAQQEVREIIKGQGKVGESDLSRLEYLKLIIKESLRLHPPGPLLIPRETIECCTIEGYKIPAKTRVFINAAAIAT
DSKVWEN-PYAFKPERFMDKTVDFKG---EDFEMLPFGAGRRGCPGMNFAVPLIGLALANLLLQFDWKLPEGMI-A
EDLEMEEAPGITVHKKIPLCLVASPNRCAAQD------------
>TliCYP8
-----------------------------MMSIFSLFPLFLFIIFLVKWFPTSSTPTKKLPPSPLKLPIIGNLHQL
G---SHPHRSLQSMSKK-YGPLMLLHFGSKPVLVASSADAAREIMKTHDLIFSTRPK-SSIPDRLLFNSKDVAFTP
YGEYWRQVRSICVIQLLSNKRVQSFRHVREEETSEMIEKIRR-----TCSSSS--INMSDVFITLTNDVVSRIALG
KK------YSHGESGRKIKALLQEFIELLGIFCVADYIPS-LAWLNRLNGLDARVERVAEQIDELMERVIDEHSKR
YDGSEEK--------GSDFVDILLEIQRTS--SGSFAVERDTLKALIMDMFAAGTDTTHSIMEWAMTELLRHPKVM
ENLQTEVRQVAQGKPEVNENEIEKMQYLKAVIKETLRLHCPVPLLIPREATQDVKVMGYDISAGTRVIVNGWAIGR
DPSFWKK-PEEFQPERFLDRDIDFRG---FNFELIPFGAGRRGCPGISFGIAVNELVLAKLMHKFNFALPEGAE-E
KDLDIPERNGNTVHRKYPLLVVATPYSM----------------
>TliCYP9
-----------------------QMVFLLNFPFSSDLALALLTLLLLRWFYALSKPQKKLPPSPRMLPIIGNLHQL
G---SYPHQSLRSLSRK-YGPLMLLHFGGAPVLVVSSADAASKIMKTHDLVFPNRPH-MSMADKLFYGSKDIVASP
YGEYWRQVRSICVLQLLSKKRVQSFQRIREEETLKMTEKIRQ-----SCSASSPVVNLSEIFAALTNDVICRVALG
RK------YSDGGNGSKFKDMLLRLGELLNAFEIADYIPW-LKWVNHFNGLDAKVERLAKEIDEFMEGVIEERRNS
KKTEAQNDGKIESI-GSDFLDILLEIQRRN--VTGFAVENDTVKALILDMFSAGTDTTYTLMEWAMAELLKHPNIM
QKLQSEIRQVAQGKPEINDDDLDKMKYLQAVIKETLRLHPPVPLLIPRQPTQDTNIMGYNIAAGTQVIVNAWAIAR
DPQLWEE-PEEFRPDRFLNSVIDFRG---SHFEFIPFGAGRRGCPGAAFAIAVNELALAKLIHKFNFALPDGKK-P
EDLDITESSGLTVHTRLPVLAVASPYGGTKSVD-----------
>SnvNO
------------------------MELLLNPSLFSLLSLLFFVIFLFKRLYASPTCQRKVPPSPPKLPVVGNLHQV
G---SLPHRSLQSLSKK-YGPLMLLHFGSVPVLVASSAEAAREIMKNHDVVFSTRPK-SNISDKLTYGSKDVAFSP
YGEYWRRVRSICVNHLLSNQRVKSFRHIMEEETRKMIENINE-----RCVSSSLPVNLSDFTITLTNSIICTMAFG
RK------HCDVENMRKIKAMLAGFEEILSVFDAGDYIPW-LAWVNRFTGLDDRLGKLAKQGDELVEGVIEEHMKR
KKAEGQRYDAADQAKGTDFMDILLDIYQGR--VPGFALDRDSVKAVILDMFTGGTDTIYTSIDWTIAELLRHPMVL
KKLHTEVREVAQGKSEITEEDLGKMAYLKVVIKETLRLHPPIPLLLPRESTQDISIMGYHISAGTQVIVNAWAIGR
DPLYWEN-AEEFRPERFMDGNMDFRG---FNFEYIPFGAGRRSCPALAFAIAVVELTIAKLVHKFDFTLPDGGK-P
EDLDMTEASGTTVHKQLPNVANVVATRN----------------
>CrT19H
--------------------------MLSSLKDFFVLLLPFFIGIAFIYKLWNFTSKKNLPPSPRRLPIIGNLHQL
S---KFPQRSLRTLSEK-YGPVMLLHFGSKPVLVISSAEAAKEVMKINDVSFADRPK-WYAAGRVLYEFKDMTFSP
YGEYWRQARSICVLQLLSNKRVQSFKGIREEEIRAMLEKINQ-----ASNNSS-IINGDEIFSTLTNDIIGRSAFG
RK------FSEEESGSKLRKVLQDLPPLLGSFNVGDFIPW-LSWVNYLNGFEKKLNQVSKDCDQYLEQVIDDTRKR
DEENGANNNGGNHG---NFVSVLLHLQKED--VKGFPSEKGFLKAIILDMIVGGTDTTHLLLHWVITELLKNKHVM
TKLQKEVREIVGRKWEITDEDKEKMKYLHAVIKEALRLHPSLPLLVPRVAREDINLMGYRVAKGTEVIINAWAIAR
DPSYWDE-AEEFKPERFLSNNFDFKG---LNFEYIPFGSGRRSCPGSSFAIPIVEHTVAHLMHKFNIELPNGVS-A
EDFDPTDAVGLVSHDQNPLSFVATPVTIF---------------
>Cr7DLH
---MELNFKSIIFLVFVSLTLYWVYRILDWVWFKPKKLEKCLREQGFKGNPYRLFLGDQYDSGKLIRQALTKPIGV
EEDVKKRIVPHILKTVGTHGKKSFMWVGRIPRVNITDPELIKEVLTKYYKFQKNHHDLDPITKLLLTGIGSLEGDP
WAKRRKIINAAFHFEKLK-LMLPAFYLSCRDMVTKWDNKVP--------EGGSAEVDVWHDIETLTGDVISRTLFG
SN---------FEEGRRIFELMKELTALTIDVIRSVYIPG---QRFLPTKRNNRMRAIDKEVRVRITEIINKKMKV
MKSGEAASAADDFLGILLECNLNEIKEQGN--NKSAGMTIGEIIGECKLFYFAGQDTTSTLLVWTMVLLSRFPEWQ
TRAREEVFQVFGNK-TPDYDGISHLKVITMILYEVLRLYTPVAELTK-VAHEATQLGKYFIPAGVQLMMPQILLHH
DPEIWGEDVMEFKPERFAEGVLKATK---SQGSFFPFSLGPRMCIGQNFALLEAKMAMSLILRRFSFELSPSYVHA
PFTLITMQPQYGAHLILHKL------------------------
>CrSLS
```

MEMDMDTIRKAIAATIFALVMAWAWRVLDWAWFTPKRIEKRLRQQGFRGNPYRFLVGDVKESGKMHQEALSKPMEF
NNDIVPRLMPHINHTINTYGRNSFTWMGRIPRIHVMEPELIKEVLTHSSKYQKNFDVHNPLVKFLLTGVGSFEGAK
WSKHRRIISPAFTLEKLK-SMLPAFAICYHDMLTKWEKIAE--------KQGSHEVDIFPTFDVLTSDVISKVAFG
ST---------YDEGGKIFRLLKELMDLTIDCMRDVYIPG---WSYLPTKRNKRMKEINKEITDMLRFIINKRMKA
LKAGEPG--EDDLLGVLLESNIQEIQKQGN--KKDGGMSINDVIEECKLFYFAGQETTGVLLTWTTILLSKHPEWQ
ERAREEVLQAFGKN-KPEFERLNHLKYVSMILYEVLRLYPPVIDLTK-IVHEDTKLGPYTIPAGTQVMLPTVMLHR
EKSIWGEDATEFNPMRFADGVANATK---NNVTYLPFSWGPRVCLGQNFALLQAKLGLAMILQRFKFDVAPSYVHA
PFTILTVQPQFGSHVIYKKLES----------------------
